# Heparan sulfates are critical regulators of the inhibitory megakaryocyte-platelet receptor G6b-B

**DOI:** 10.1101/584698

**Authors:** Timo Vögtle, Sumana Sharma, Jun Mori, Zoltan Nagy, Daniela Semeniak, Mitchell J. Geer, Christopher W. Smith, Jordan Lane, Scott Pollack, Riitta Lassila, Annukka Jouppila, Alastair J. Barr, Derek J. Ogg, Tina D. Howard, Helen J. McMiken, Juli Warwicker, Catherine Geh, Rachel Rowlinson, W. Mark Abbott, Harald Schulze, Gavin J. Wright, Alexandra Mazharian, Klaus Fütterer, Sundaresan Rajesh, Michael R. Douglas, Yotis A. Senis

## Abstract

The immunoreceptor tyrosine-based inhibition motif (ITIM)-containing receptor G6b-B is critical for platelet production and activation, loss of which results in severe macrothrombocytopenia and aberrant platelet function in mice and humans. Using a combination of immunohistochemistry, affinity chromatography and proteomics, we identified the extracellular matrix heparan sulfate (HS) proteoglycan perlecan as a G6b-B binding partner. Subsequent *in vitro* biochemical studies and a cell-based genetic screen demonstrated that the interaction is specifically mediated by the HS chains of perlecan. Biophysical analysis revealed that heparin forms a high-affinity complex with G6b-B and mediates dimerization. Using platelets from humans and genetically-modified mice, we demonstrate that binding of G6b-B to HS and multivalent heparin inhibits platelet and megakaryocyte function by inducing downstream signaling via the tyrosine phosphatases Shp1 and Shp2. Our findings provide novel insights into how G6b-B is regulated and contribute to our understanding of the interaction of megakaryocytes and platelets with glycans.

## INTRODUCTION

Platelets are highly reactive anucleated cell fragments, produced by megakaryocytes (MKs) in the bone marrow, spleen and lungs. In an intact vasculature, platelets circulate in the blood stream for about eight to ten days and are finally cleared by reticulo-endothelial system. Upon vascular injury, however, platelets adhere to the exposed vascular extracellular matrix (ECM), become activated and form a hemostatic plug that seals the wound. Platelet activation must be tightly regulated to avoid hyperactivity and indiscriminate vessel occlusion (Bye, Unsworth, & Gibbins, 2016; Jackson, 2011). The mechanisms that inhibit platelet activation include extrinsic factors, such as endothelial-derived nitric oxide and prostacyclin, and intrinsic factors, such as immunoreceptor tyrosine-based inhibition motif (ITIM)-containing receptors (Coxon, Geer, & Senis, 2017; Nagy & Smolenski, 2018).

G6b-B is a unique platelet ITIM-containing receptor that is highly expressed in mature MKs and platelets (Coxon et al., 2017; Senis et al., 2007). It is a type I transmembrane protein that consists of a single N-glycosylated immunoglobulin-variable (IgV)-like domain in its extracellular region, a single transmembrane domain and a cytoplasmic tail containing an ITIM and an immunoreceptor tyrosine-based switch motif (ITSM). The central tyrosine residue embedded in the consensus sequence of the ITIM ([I/V/L]xYxx[V/L]) become phosphorylated by Src family kinases (SFKs) and subsequently acts as a docking site for the SH2 domain-containing protein-tyrosine phosphatases (Shp)1 and 2 (Mazharian et al., 2012; Senis et al., 2007). The canonical mode of action of ITIM-containing receptors is to position these phosphatases in close proximity to ITAM-containing receptors, allowing them to dephosphorylate key components of the ITAM signaling pathway and attenuate activation signals. The inhibitory function of G6b-B has been demonstrated in a heterologous cell system, by antibody-mediated crosslinking of the receptor in platelets and G6b-B knockout (*KO*) mouse models (Mazharian et al., 2012; Mori et al., 2008; Newland et al., 2007). Findings from these mice demonstrated that the function of G6b-B goes beyond inhibiting signaling from ITAM-containing receptors (Mazharian et al., 2013; Mazharian et al., 2012). These mice develop a severe macrothrombocytopenia and aberrant platelet function, establishing G6b-B as a critical regulator of platelet activation and production. This phenotype was also observed in a G6b-B loss-of-function mouse model (*G6b* diY/F) in which the tyrosine residues within the ITIM and ITSM were mutated to phenylalanine residues, abrogating binding of Shp1 and Shp2 to G6b-B and downstream signaling (Geer et al., 2018). Moreover, expression of human G6b-B in mouse platelets rescued the phenotype of G6b-B-deficient mice, demonstrating that human and mouse G6b-B exert the same physiological functions (Hofmann et al., 2018). Importantly, null and loss-of-function mutations in human G6b-B have been reported to recapitulate key features of the *G6b* KO and loss-of-function mouse phenotypes, including a severe macrothrombocytopenia, MK clusters in the bone marrow and myelofibrosis (Hofmann et al., 2018; Melhem et al., 2016). Despite the vital role of G6b-B in regulating platelet production and function, its physiological ligand was not known. Although a previous study demonstrated that G6b-B binds to the anti-coagulant heparin, however, the functional significance and physiological relevance of this finding have proved elusive (de Vet, Newland, Lyons, Aguado, & Campbell, 2005).

Proteoglycans comprise a heterogeneous family of macromolecules, consisting of a core protein and associated unbranched glycosaminoglycan (GAG) side-chains. Heparan sulfates (HS) are a specific subgroup of GAGs, defined by their basic disaccharide unit. They are structurally-related to heparin, which is produced as a macromolecular proteoglycan by tissue-resident mast cells (Lassila, Lindstedt, & Kovanen, 1997) and, following chemical or enzymatic processing, serves as an anti-coagulant (Chandarajoti, Liu, & Pawlinski, 2016; Meneghetti et al., 2015). One of the best studied HS proteoglycan is perlecan, which is synthesized and secreted by endothelial and smooth muscle cells into the vessel wall. It is comprised of a large 400 kDa core protein and has three HS chains attached to its N-terminus. A number of proteins reportedly interact with the HS chains and protein core of perlecan, among them are structural components of the ECM, including laminin, collagen IV and fibronectin, and fibroblast growth factor-2 (Nugent, Nugent, Iozzo, Sanchack, & Edelman, 2000; Whitelock, Melrose, & Iozzo, 2008). Of note, the proteolytically released C-terminal fragment of perlecan, called endorepellin, binds to integrin α2β1 and enhances collagen-mediated platelet activation (Bix et al., 2007). Perlecan has also been shown to exert anti-thrombotic properties in an ovine vascular graft model through its HS side-chains, however the underlying mechanism has not been defined (Lord et al., 2009).

In this study, we identified the physiological ligand of G6b-B, the molecular basis of the G6b-B ligand interactions and the mechanism underlying physiological effects. Our findings demonstrate that G6b-B binds the HS chains of perlecan, as well as to heparin, eliciting functional responses in MKs and platelets. Moreover, we also show that a cross-linked, semisynthetic form of heparin, called anti-platelet anti-coagulant (APAC) (Lassila & Jouppila, 2014), beyond inhibiting collagen-mediated platelet aggregation, induces robust phosphorylation and downstream signaling of G6b-B. Collectively, these results reveal that HSs regulate G6b-B signaling and function, providing a novel mechanism by which MK and platelet function is regulated.

## RESULTS

### Identification of perlecan as a ligand of G6b-B

To identify the tissue expressing the physiological ligand of G6b-B, we generated a recombinant G6b-B Fc-fusion protein (mG6b-B-Fc), consisting of the murine G6b-B ectodomain and the human IgG-Fc tail, to mediate dimer formation, that we used to stain frozen mouse tissue sections. We consistently observed prominent staining in large vessels, including the vena cava and aorta, and also in smaller vessels in liver and spleen, not observed with the negative control (IgG-Fc) (Figure 1), suggesting the presence of G6b-B ligand in the vessel wall.

**Figure 1.**
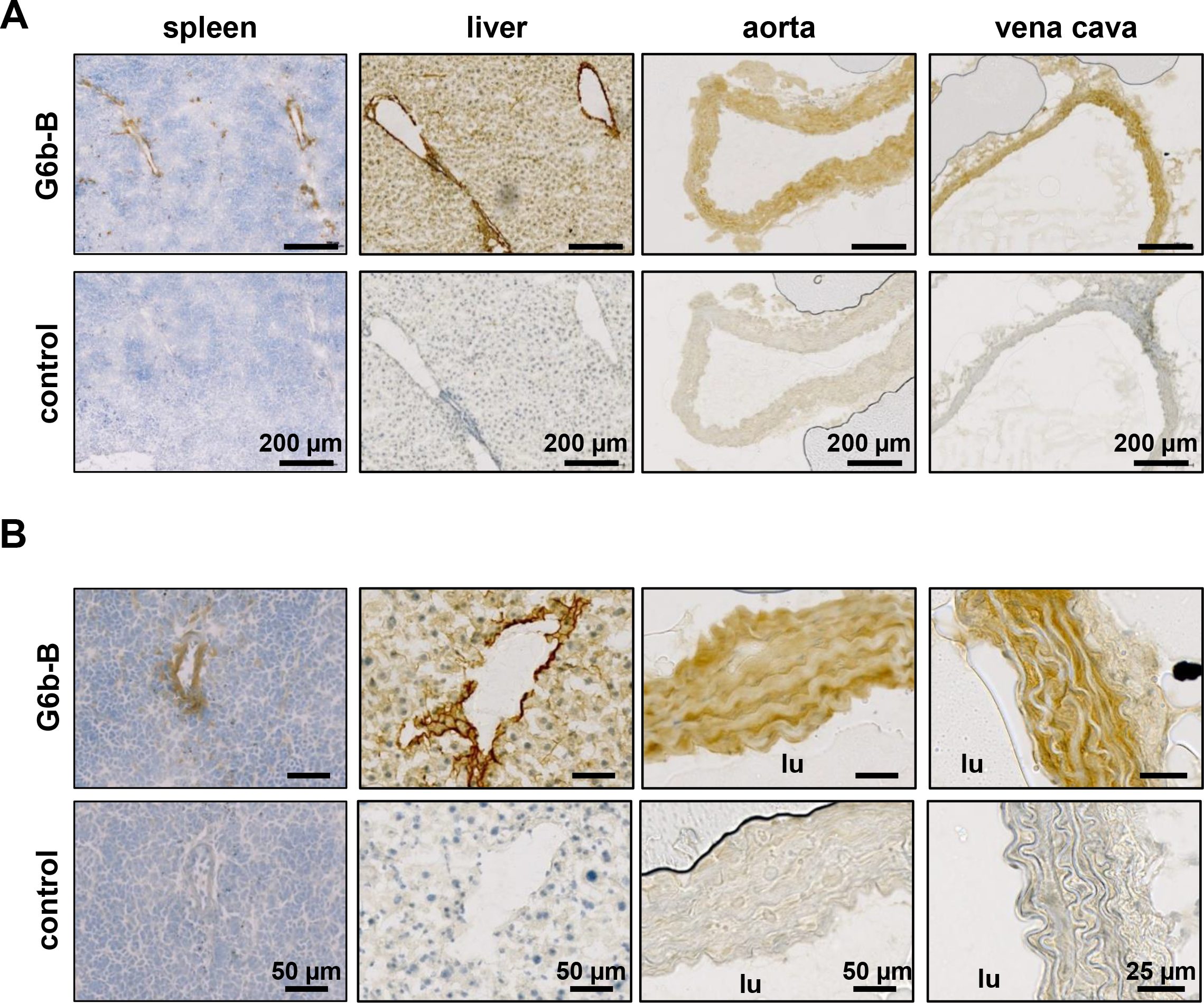
Prominent binding of mG6b-B-Fc to the vessel wall. Immunohistochemistry staining of frozen mouse tissue sections with mG6b-B-Fc or human IgG-Fc fragment (control). Bound protein was visualized using a secondary anti-human-Fc-HRP antibody and DAB substrate, prior to counterstaining with hematoxylin. The images were captured by a Zeiss Axio Scan.Z1 slidescanner, and images were exported with the Zeiss Zen software. **(A)** Overview and **(B)** zoom-in images for the indicated tissues are shown. lu, vessel lumen

To identify the identity of the ligand we incubated vena cava homogenates with mG6b-B-Fc and protein G sepharose beads to precipitate G6b-B binding partners. SDS-PAGE and Colloidal Coomassie staining revealed bands of high molecular weight that were absent in the negative control (IgG-Fc pulldown, data not shown). Bands were excised and proteins identified by mass spectrometry, revealing basal membrane-specific HS proteoglycan (HSPG) core protein or perlecan, as the most abundant protein specifically pulled-down with mG6b-B-Fc (Table 1).

**Table 1.**
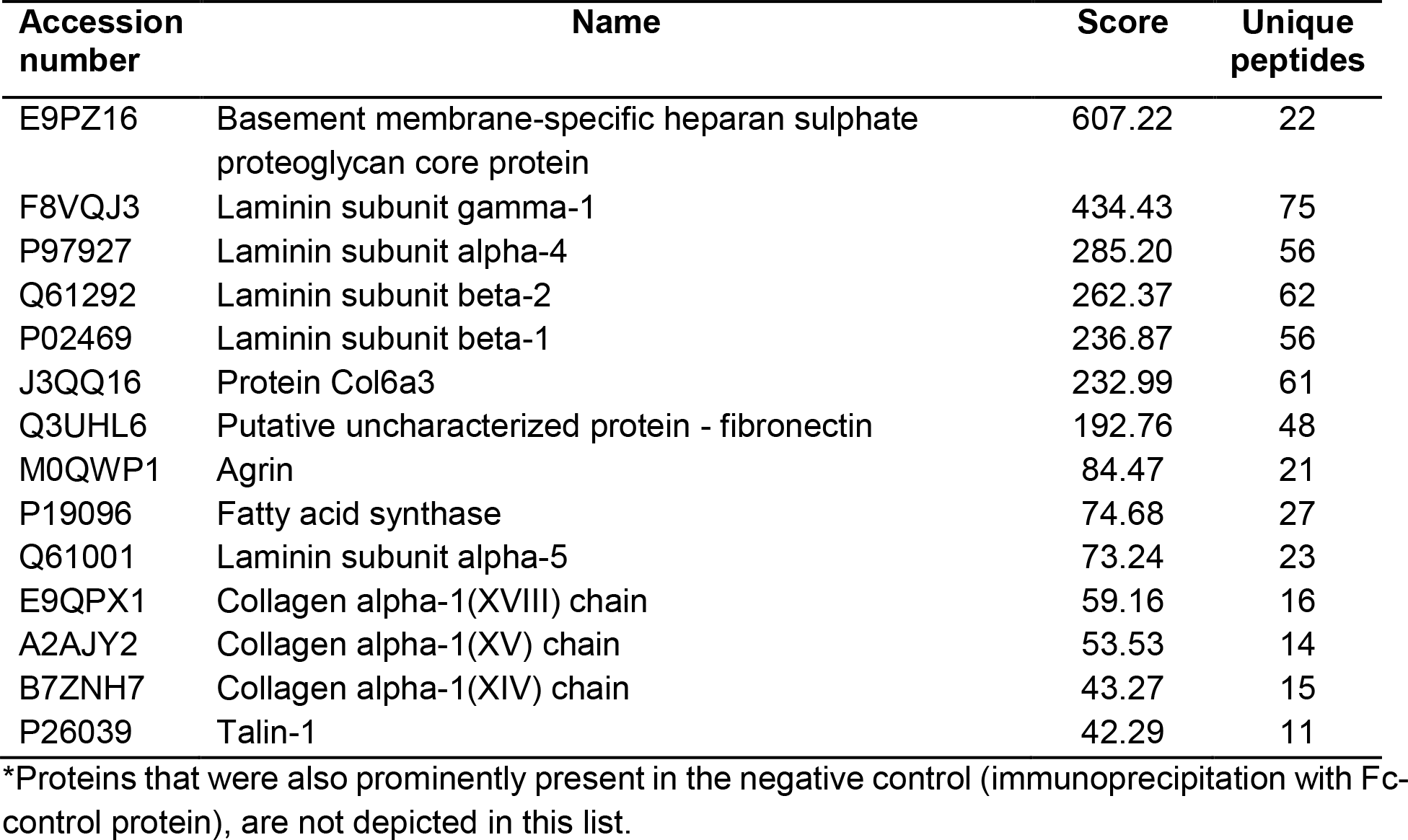
List of proteins immunoprecipitated with mG6b-B-Fc from vena cava lysates*

The interaction with perlecan was verified using an *in vitro* binding assay, measuring binding of soluble mG6b-B-Fc to immobilized molecules. mG6b-B-Fc bound robustly to perlecan, but not to BSA (control) or other ECM molecules, including collagen I and IV, various forms of laminin (111, 411, 421, 511 and 521), fibronectin or the related and recombinantly expressed HSPGs syndecan-2 or agrin (Figure 2A). Hence, laminin and collagen identified by G6b-B pulldown and mass spectrometry (Table 1) most likely represented perlecan-associated proteins (Battaglia, Mayer, Aumailley, & Timpl, 1992), rather than direct binding partners of G6b-B. Human G6b-B-Fc (huG6b-B-Fc) showed similar binding characteristics as mG6b-B-Fc (Figure 2A).

**Figure 2.**
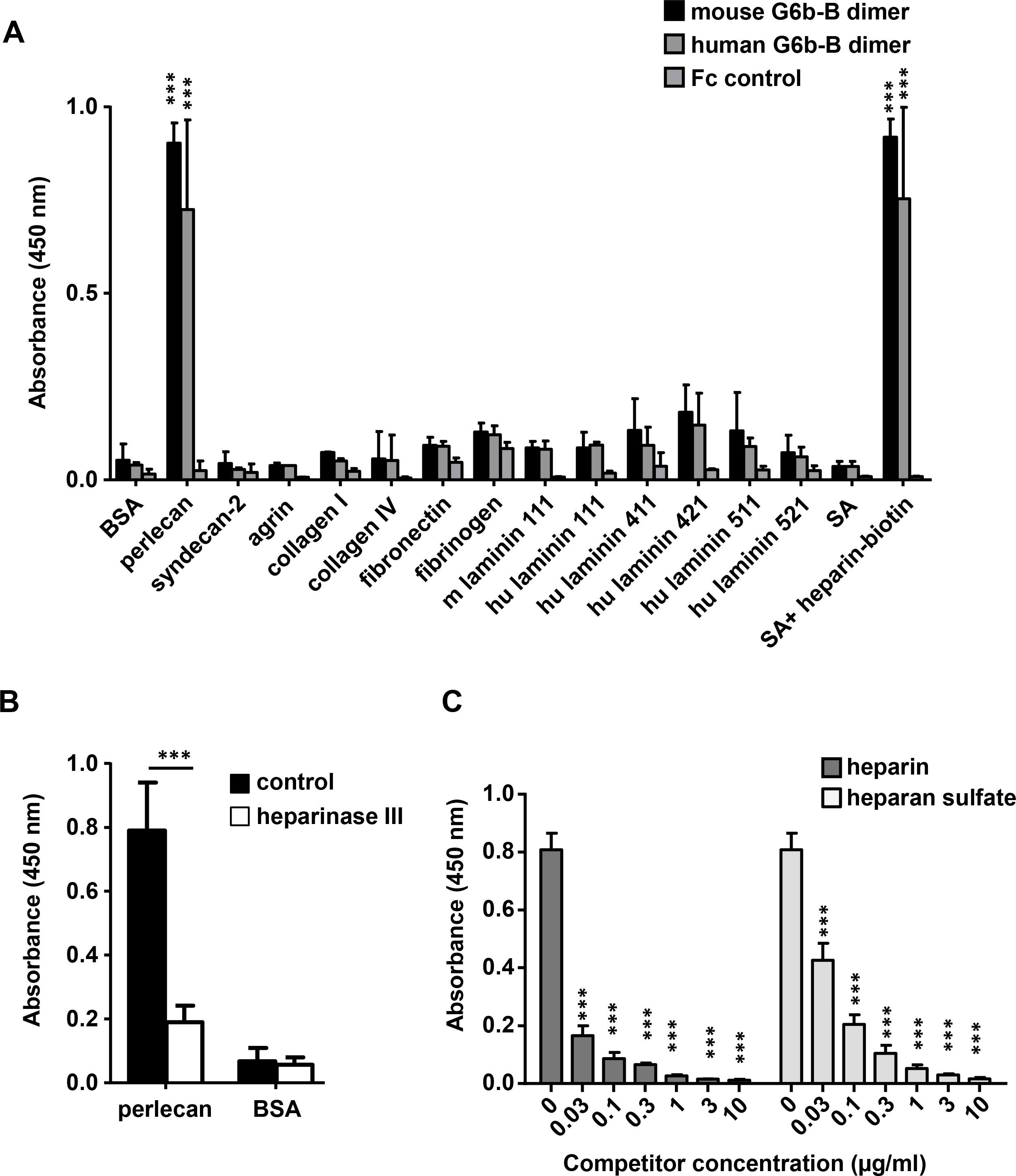
G6b-B-Fc binds to heparan sulfate side-chains of perlecan. **(A)** 96-well plates were coated with the indicated substrates (5 μg/ml) and incubated with mouse G6b-B-Fc (10 μg/ml), human G6b-B-Fc (30 μg/ml) or Fc-control (10 μg/ml). Bound protein was detected with an anti-human-Fc-HRP antibody and TMB substrate. n=2-4; SA, streptavidin; **(B)** Perlecan and bovine serum albumin (BSA) were treated or not with heparinase III (5 mU/ml) prior to blocking and mG6b-B-Fc binding was measured. n=4; **(C)** mG6b-B binding to immobilized perlecan was measured in the presence of the indicated concentrations of heparin and heparan sulfate; n=3; P-values were calculated using ordinary one-way ANOVA with Dunnett’s post-hoc test and asterisks denote statistical significance compared to respective control. ***, *P<0.001*

Treatment of perlecan with the enzyme heparinase III, which removes the HS side-chains, significantly reduced G6b-B binding to immobilized perlecan (Figure 2B), indicating binding of G6b-B to the HS side-chains rather than the protein core. This observation was further supported by a competition assay, in which the addition of soluble HS inhibited the binding of G6b-B to immobilized perlecan (Figure 2C). Of note, unfractionated heparin, which is closely related to HS, also interfered with G6b-B binding to perlecan and streptavidin-immobilized biotin-conjugated heparin directly bound to G6b-B-Fc (Figure 2A).

To gain further insights into the structural requirements of the G6b-B ligand interaction, we tested heparin oligomers of different length (4, 8, 12 and 20 saccharide units, dp4, dp8, dp12 and dp20, respectively) and selectively desulfated heparin molecules for their binding to G6b-B. In a competition assay, only oligomers of at least 8 saccharides were able to partially block binding of G6b-B to heparin-biotin, suggesting this to be the minimum length required for this interaction (Figure 2–figure supplement 1A). In addition, high sulfation of the glycan was found to be important for G6b-B binding, since a loss in one sulfation site resulted in a significant drop in the ability to block G6b-B binding to native heparin (Figure 2–figure supplement 1B).

Since that the binding assay results suggested that the G6b-B ligand was primarily composed of HS glycans, we opted to confirm and extend these finding by using a genome-scale cell-based CRISPR knockout (KO) screening approach, identifying all genes required for the synthesis and cell surface display of the G6b-B ligand (Sharma, Bartholdson, Couch, Yusa, & Wright, 2018). We observed that a fluorescently labelled highly-avid recombinant G6b-B binding reagent robustly stained several human cell lines providing the basis for a cellular genetic screen (Figure 3A). A genome-wide mutant cell library was generated by transducing Cas9-expressing HEK293 cells with a library of lentiviruses each encoding a single gRNA from a pool of 90,709 individual gRNAs targeting 18,009 human genes (Sharma et al., 2018). Transduced cells that had lost the ability to bind to the recombinant protein were isolated using fluorescent-activated cell sorting and genes required for cell surface binding of G6b-B were identified by comparing the relative abundance of gRNAs in the sorted versus unsorted control population (Li et al., 2014). Using this strategy, we unambiguously identified many genes required for HS biosynthesis, beginning with the generation of the tetrasaccharide linkage on the serine residue of the protein backbone (*B3GAT3*, *XYLT2*, *B4GALT7*), the commitment towards the HS pathway (*EXTL3*), HS chain polymerization (*EXT1/2*), and HS chain modification (*NDST1*, *HS2ST1*) (Figure 3B). Of particular note, genes encoding the enzymes chondroitin sulfate N-acetylgalactosaminyltransferase 1 and 2 (*CSGALNACT1/2*), which are essential for the commitment towards the biosynthesis of chondroitin sulfate chains were not identified, demonstrating that G6b-B binding to HEK293 cells is mediated by HS, but not by chondroitin sulfate (Figure 3B). Moreover, addition of heparin, but not chondroitin sulfate, inhibited G6b-B binding to HEK293 cells (Figure 3C). We also identified *SLC35B2* (Solute Carrier Family 35 Member B2), a gene encoding a transporter protein that translocates 3’-phosphoadenosine-5’-phosphosulfate, from the cytosol into the Golgi apparatus, where it is used as a sulfate donor for sulfation of glycoproteins, proteoglycans, and glycolipids. We validated the involvement of sulfated HSs in mediating G6b-B binding to cells by individually targeting *SLC35B2* and were able to demonstrate that this led to a loss in G6b-B binding relative to the parental cell line (Figure 3D). Together, this genetic screen provides further evidence, corroborating our *in vitro* binding data, that the physiological ligand of G6b-B is negatively charged HS.

**Figure 3.**
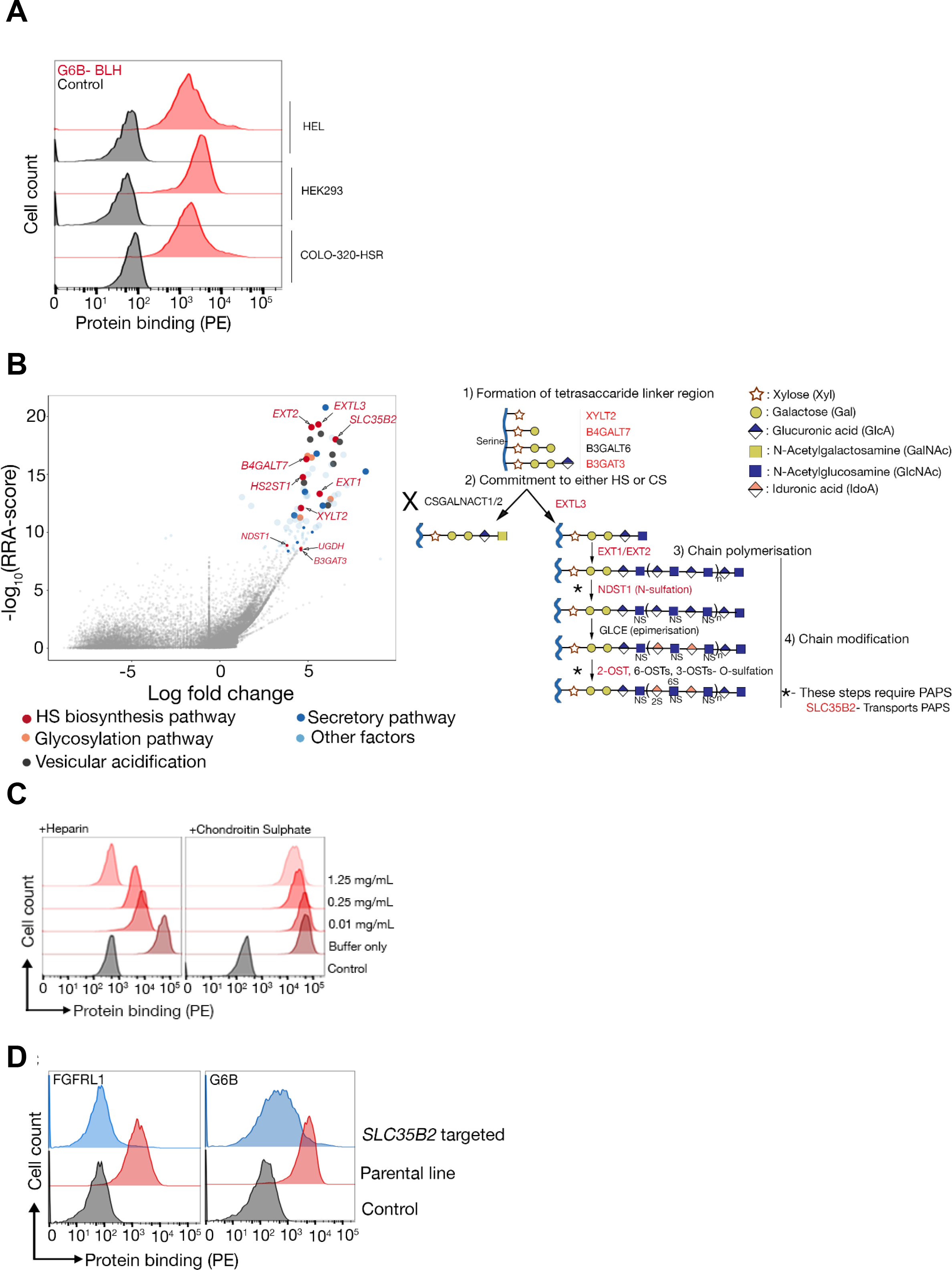
The heparan sulfate biosynthesis pathway is required for G6b-B binding to HEK293 cells. **(A)** Recombinant G6b-B - produced as a monomeric biotinylated protein and conjugated to streptavidin-PE to generate an avid probe - binds to HEL, HEK293 and COLO-320-HSR cells. **(B)** A genome-wide loss-of-function approach identifies the HS biosynthesis pathway as the required factor for mediating binding of recombinant G6b-B to HEK293 cell (left panel). X- and y-axis represent log-fold-change (LFC) and Robust Rank Aggregation (RRA) score calculated using the MAGeCK software, respectively. Circles represent individual genes and sizes represent the false-discovery rate: large circle = FDR<1%, small circle = 1% < FDR < 5%. Genes with FDR < 5% are color coded according to their functional annotation and genes corresponding to the HS biosynthesis pathway are additionally named. The HS biosynthesis pathway is depicted in the right panel with the genes identified in the loss-of-function approach highlighted. Similar results were obtained in HEL cells (not shown). **(C)** G6b-B binding to HEK293 cells was measured by flow cytometry in the presence or absence of the indicated concentration of heparin or chondroitin sulfate. One representative out of three experiments is shown. **(D)** G6b-B loses its binding to cell lines where SLC35B2, required for the sulfation of GAGs, is targeted. To ensure that the KO cells lack GAGs, a known HS binding protein is used as a control, which loses binding on these cell lines.

### Molecular basis of G6b-B interaction with HS side-chains of perlecan

The extracellular domain of G6b-B is enriched in positively charged residues, especially arginines (12 in 125 amino acids; 9.6% vs 5.6% average frequency in mammalian membrane proteins (Gaur, 2014)) which are known to mediate strong binding to heparin (Margalit, Fischer, & Ben-Sasson, 1993). Prior to obtaining the crystal structure, we generated a structural model of G6b-B using template-based tertiary structure prediction (RaptorX Structure Prediction server) and used this model to aid identification of candidate residues for mutagenesis. Examination of the model showed four basic residues (Lys54, Lys58, Arg60, Arg61) in close spatial proximity to each other on a solvent-exposed loop. We tested whether these amino acids are involved in heparin binding, by generating a mutant G6b-B (K54D, K58D, R60E, R61E; Figure 4–figure supplement 1A) and comparing heparin binding to WT G6b-B in transiently transfected CHO cells. An anti-G6b-B monoclonal antibody demonstrated robust cell surface expression of mutant G6b-B that was comparable to that of WT G6b-B, suggesting the quadruple mutation did not disrupt protein folding or expression (Figure 4–figure supplement 1B). Cells expressing WT G6b-B showed an increase in heparin binding, as compared to non-transfected cells, while the cells expressing mutant G6b-B showed minimal binding compared to controls, demonstrating that these amino acids (or a subset thereof) are required for ligand binding (Figure 4–figure supplement 1C).

**Figure 4.**
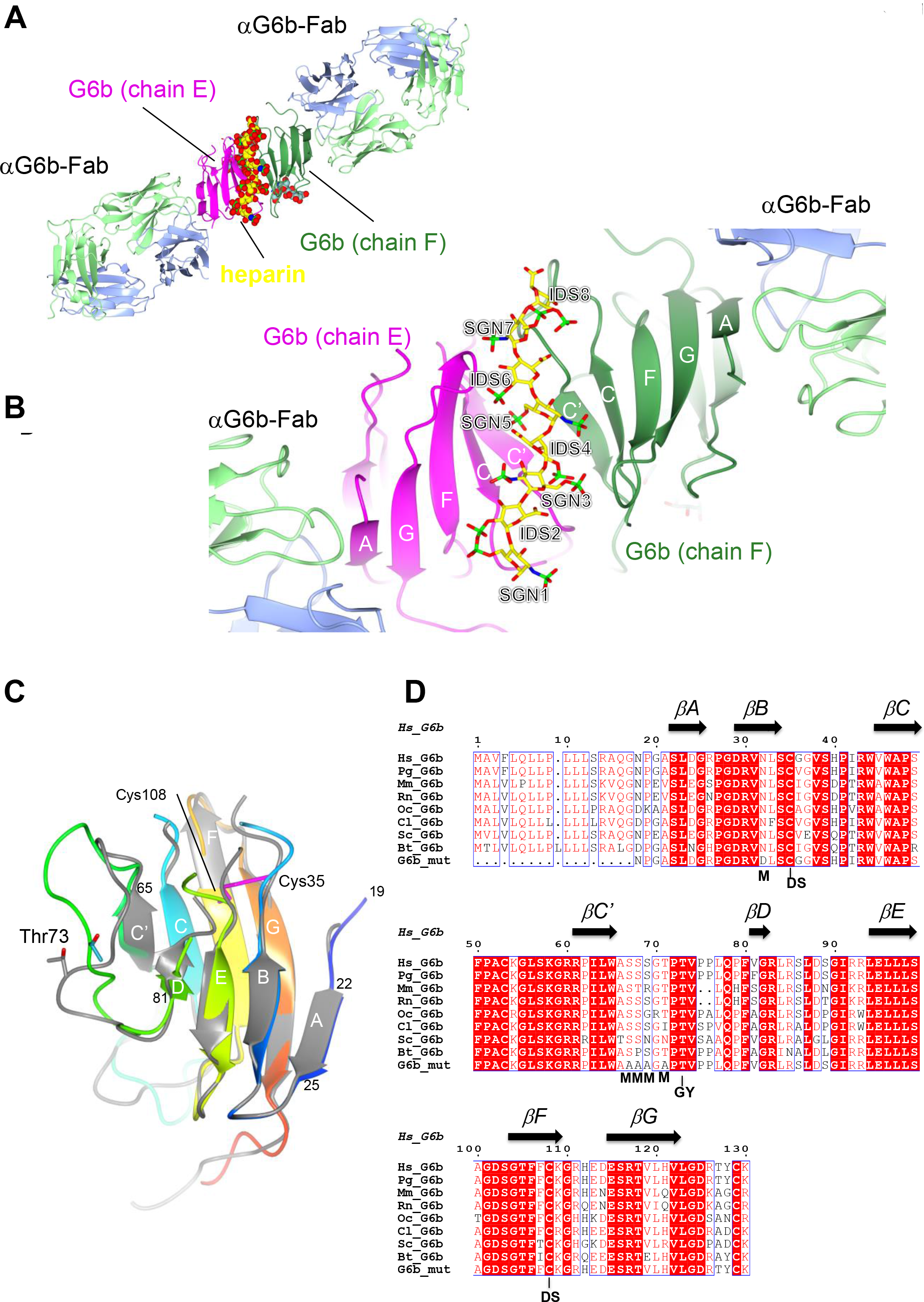
Ribbon representation of the ternary complex of the extracellular domain (ECD) of human G6b-B bound to heparin and the Fab fragment of a G6b-B-specific antibody. **(A)** Overview of the structure with G6b-B colored in magenta and dark green, heparin shown as spheres, and the Fab fragment chains in light green / light blue, respectively. The assembly represents the asymmetric unit of the crystal lattice (space group *C*2). (B) Close-up view illustrating the position of the heparin ligand relative to the secondary structure of the G6b-B dimer. Heparin residues (shown as sticks) are sulfated D-glucosamine (SGN) and L-iduronic acid (IDS). The color coding of heparin atoms is C – yellow, O – red, N – blue, S – green and β-strands in G6b-B are labelled according to the canonical Ig-fold. **(C)** Superposition of chains F (rainbow colored) and E (grey) of the G6b-B ECD. Strands are labelled according to the canonical β-sandwich topology of the variable Ig domain. The fold of G6b-B deviates from the canonical Ig fold in missing strand C”, and in strand A being part of the β-sheet of strands B, E and D. Chain F is color ramped from blue (N-terminus) to red (C-terminus), and the position of the disulfide bond (Cys35–Cys108) is indicated by sticks in magenta. The glycosylation site Thr73 is shown (sticks) with glycosyl groups omitted from the view. **(D)** Multiple sequence alignment of G6b-B orthologues across mammalian species with secondary structure elements indicated above the sequence. Residue numbers refer to the sequence of human G6b. Conserved residues are boxed, with identities shown as white letters on red background. Species abbreviations are Hs – Homo sapiens, Pg – Pan troglodytes (chimpanzee), Mm – Mus musculus (mouse), Rn – Rattus norvegicus (rat), Oc – Oryctolagus cuniculus (rabbit), Cl – Canis lupus familiaris (dog), Sc – Sos scroftus (wild boar), Bt – Bos taurus (cattle). G6b_mut is the sequence of the recombinant human G6b-B ECD used in crystallization, with mutations of the 5 putative glycosylation sites (marked with M). GY indicates the retained O-glycosylation site and DS indicates the disulfide cysteine residues

### The crystal structure of the G6b-B ECD-dp12-Fab complex

Subsequent to the tertiary structure prediction, we were able to generate crystals of the ternary complex of the ectodomain of G6b-B bound to the heparin oligosaccharide dp12, scaffolded by a G6b-B-specific Fab fragment and determined the structure of this complex by X-ray crystallography to 3.1 Å resolution (Figure 4 and Table 2). The complex encompasses 6 protein subunits, a dimer of G6b-B and two Fab fragments. As expected for a Fab-scaffolded structure, crystal packing contacts occur predominantly between the Fab fragment subunits (Figure 4– figure supplement 2A), but sparse direct contacts between symmetry-related G6b-B subunits also occurred (Figure 4–figure supplement 2B).

**Table 2.** Crystallographic data collection and refinement statistics for the G6b-B ECD-dp12-Fab complex.

| **X-ray diffraction data** |  |
| --- | --- |
| Beamline | I03, Diamond Light Source |
| Wavelength (Å) | 0.97624 |
| Space group | C2 |
| Cell parameters (Å) | 183.8, 72.34, 131.0, $\beta = 124.5^\circ$ |
| Complexes per asymmetric unit | 1 |
| Resolution range (Å) | 65.27 – 3.13 |
| High resolution shell (Å) | 3.18 – 3.13 |
| Rmerge (%)<sup>1)</sup> | 17.0 (146.6) |
| Total observations,<br>unique reflections | 74,255 / 24,543 |
| I/$\sigma$(I)<sup>1)</sup> | 4.0 (0.7) |
| Completeness (%)<sup>1)</sup> | 97.2 (98.2) |
| Multiplicity<sup>1)</sup> | 3.0 (3.1) |
| CC<sub>1/2</sub><sup>1), 2)</sup> | 0.991 (0.348) |
| **Refinement** |  |
| Resolution range | 63.1 – 3.13 |
| Unique reflections | 24543 |
| R<sub>cryst</sub>, R<sub>free</sub> (%) | 22.6, 26.0 |
| No of non-H atoms | 7,852 |
| RMSD bonds (Å) | 0.01 |
| RMSD angles ($^\circ$) | 1.18 |
| B-factors |  |
| Wilson (Å<sup>2</sup>) | 77.5 |
| Average overall (Å<sup>2</sup>) | 84.7 |
| RMSD B-factors (Å<sup>2</sup>) | 5.737 |
| Ramachandran statistics<sup>3)</sup> |  |
| Favoured regions (%) | 91.2 |
| Allowed regions (%) | 8.3 |
| Disallowed (%) | 0.5 |

1) parentheses refer to the high resolution shell
2) as defined in (Karplus & Diederichs, 2012)
3) calculated using molprobity (Williams et al., 2018)

Confirming the fold of the predicted model, the ectodomain of G6b-B forms an immunoglobulin-like fold of a topology closely resembling the structure of a variable immunoglobulin (Ig) domain (Figure 4C) (Brändén & Tooze, 2009). A disulfide bond between cysteine residues 35 and 108 (strands B and F, respectively) stabilizes the immunoglobulin (Ig) fold (Figure 4C). The backbone does not form the canonical strand C”, and only a very short strand D. In a canonical Ig domain, strand A is part of the sheet formed by strands B–E–D, but in the case of G6b-B, it is part of the opposite sheet (strands C’–C–F–G). The two G6b-B subunits (peptide chains E and F in the coordinate set) superimpose closely relative to the core β-sandwich structure, but divert markedly from each other in the loop connecting strands C’ and D (residues 66 to 81, Figure 4C). This loop includes several putative O-glycosylation sites (Figure 4D), which were mutated to Ala to ensure homogenous glycosylation of the protein. However, the O-linked glycosylation site Thr73 was retained, and electron density shows the presence of three saccharides attached to Thr73 in both peptide chains (Figure 4 –figure supplement 3). Although the electron density (resolution 3.1 Å) does not allow one to identify the saccharides unequivocally, the groups could be modelled as galactose, α-N-acetyl-D-galactosamine and O-sialic acid, respectively. These glycosyl groups are well separated from the heparin oligosaccharide.

The ectodomain of G6b-B assembles into an apparent dimer with a pseudo two-fold symmetry oriented perpendicular to the extended β-sheet that forms the heparin binding site (Figure 4C). Dimer formation of G6b-B is driven by the heparin ligand, as demonstrated by size exclusion chromatography (Figure 5). The interface between chains E and F buries approximately 800 Å^2^ of solvent accessible surface area. In line with the modest surface area buried between the two subunits, the interface analysis using the PISA software does not predict a stable complex (Krissinel & Henrick, 2007), consistent with the observation that ectodomain dimerization is induced by the heparin ligand. Non-covalent contacts between the two chains consist almost entirely of van der Waals (vdW) and hydrophobic interactions, with Trp65^F^ and Pro62^F^ positioned centrally in the interface, contacting Pro62^E^ and Arg61^E^, while Trp65^E^ forms vdW contacts with Val77^F^. There are very few H-bond interactions (Ser57^E^-Oγ – Ala66^F^-O/Ala68^F^-N; Lys58^E^-Nζ – Arg43^F^-O) across the interface, and notably the central β-sheet (strands C’–C–F– G–A) is not continuous in that it lacks main chain-main chain hydrogen bonds between the C’ strands of opposing protomers (Figure 4B). Nevertheless, dimerization creates a deep cleft, into which the heparin ligand inserts (Figure 6A). Crystallization involved a dodeca-saccharide, of which 8 residues are visible in the electron density map (Figure 6–figure supplement 1), with the central residues 4 and 5 representing sulfated D-glucosamine (SGN) and L-iduronic acid (IDS), respectively. While the ligand binding cleft provides partial charge complementarity to the sulfate groups of the heparin ligand (Figure 6A), perhaps surprisingly only one sulfate group (residue SGN5) forms ionic interactions with basic side-chains (SGN5-O2S – Arg60^F^-Nε3.3 Å, SGN5-O3S– Lys109^F^-Nζ 3.2 Å, superscript refers to the chain ID, Figure 6B, C). The other 8 polar contacts (within a distance cut-off of 4 Å) involving sulfate groups are with backbone amides (Arg60^E^, Glu113^E^, His112^E^, 2.8 – 3.3 Å) rather than side-chains, while 9 residues, including Lys109^E^, form vdW interactions with the ligand (Figure 6B, C). There is exquisite shape complementarity between the heparin and the surface of the G6b-B dimer, even though the S-shaped ligand does only partially fill the ligand binding cleft.

**Figure 5.**
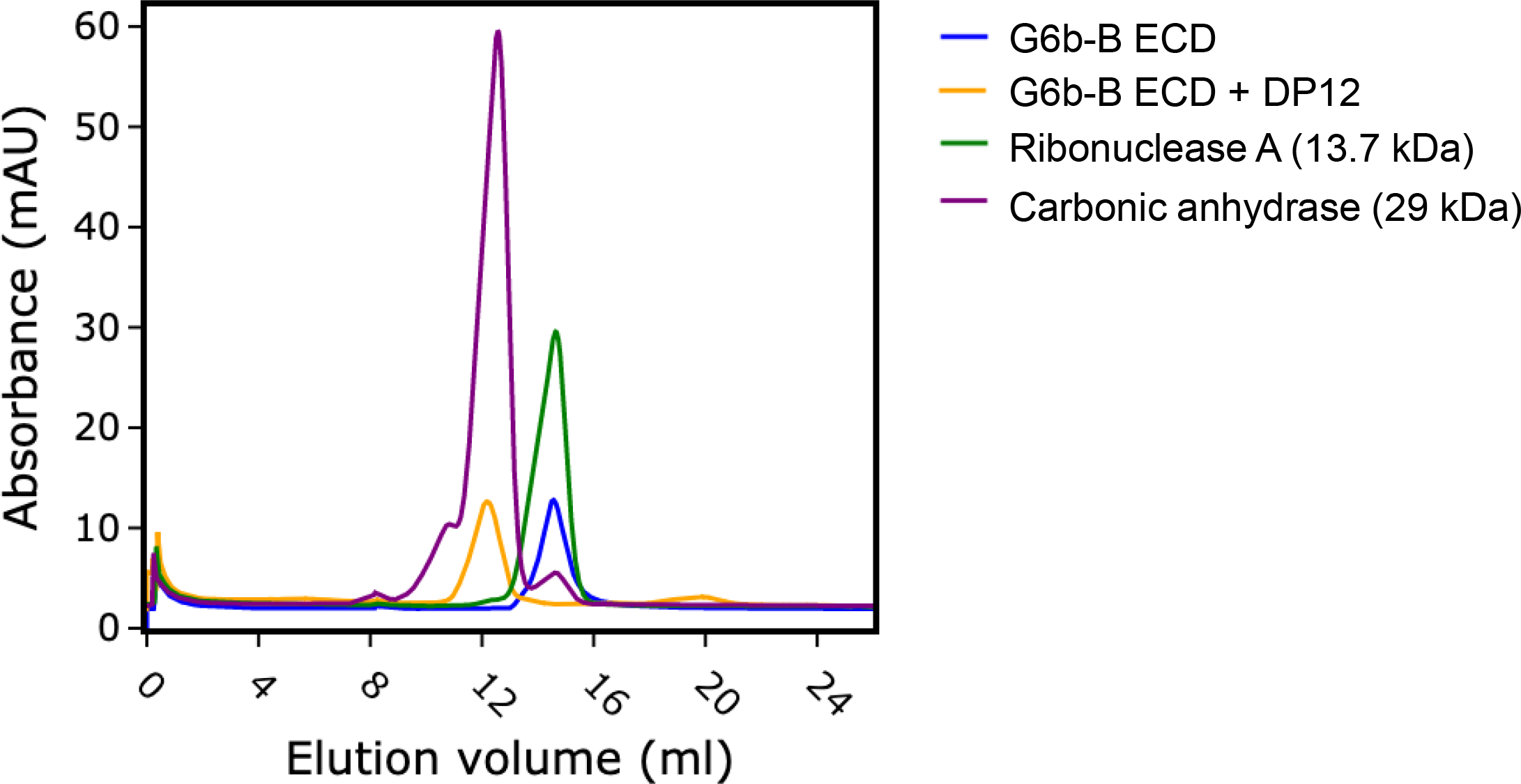
Heparin induces G6b-B dimer formation. Size exclusion chromatography of G6b-B ECD. Protein was either analyzed immediately or incubated at 4°C for 1.5 hours in the presence of dp12 before analysis on a Superdex 75 10/300 GL column. Molecular weights were estimated using a calibration curve. Values of 30.8kDa and 12.9 kDa were obtained for G6b-B ECD in the presence and absence of dp12 (approx. 3.6 kDa), respectively. Ribonuclease A (13.7 kDa) and carbonic anhydrase (29 kDa) are shown for comparison.

**Figure 6.**
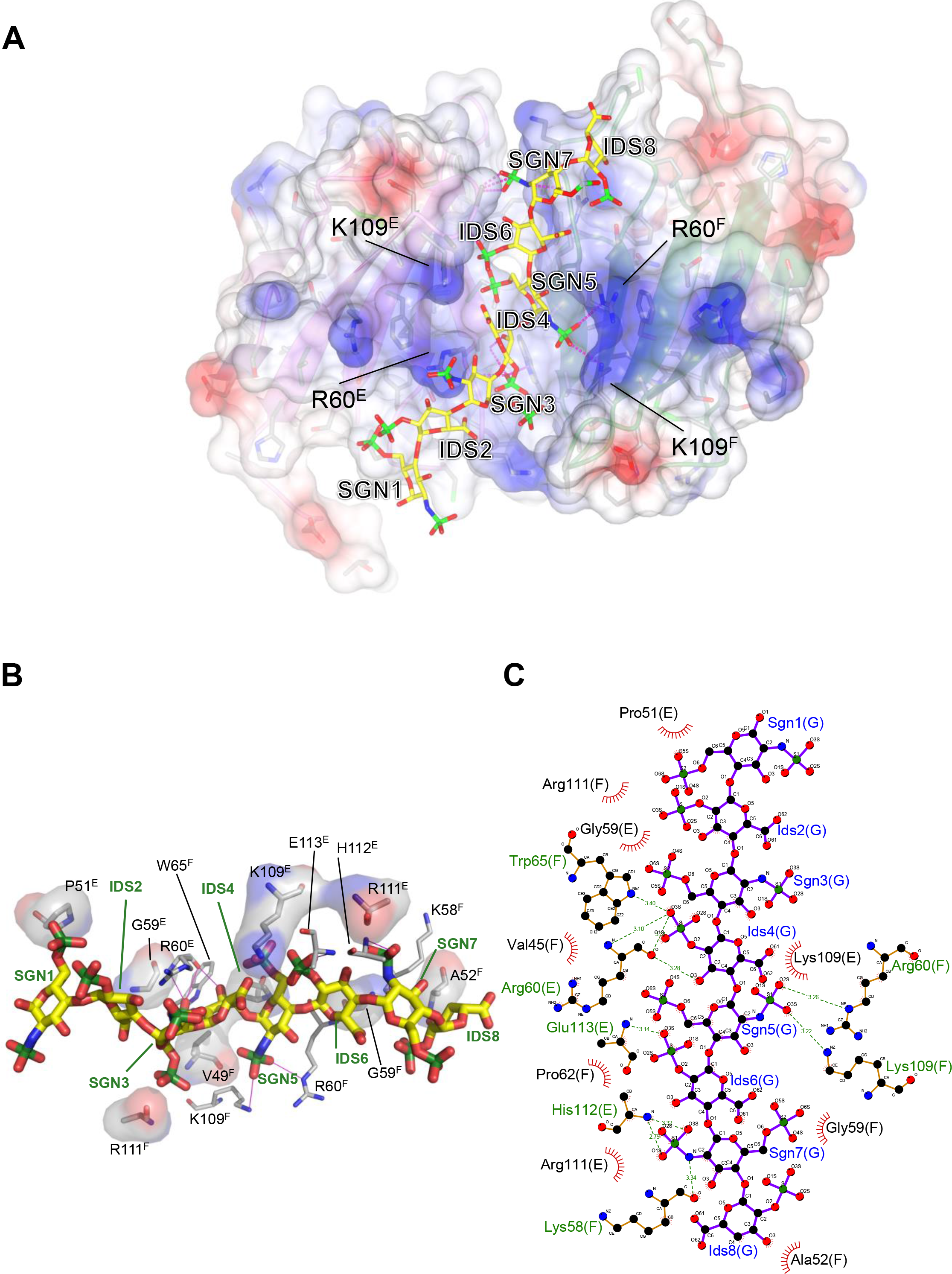
Electrostatic surface potential of the G6b-B ECD and representation of non-covalent contacts between heparin and G6b. **(A)** The G6b-B dimer is shown with a translucent surface colored according to electrostatic surface potential (calculated using CCP4mg). The heparin ligand is shown as a stick model and polar contacts are indicated by dashed lines in magenta. Selected residues are labelled with superscripts indicating the relevant G6b-B protein chain. **(B, C)** Representation of non-covalent contacts between heparin and the G6b dimer. **(B)** Residues of G6b-B forming non-covalent contacts with heparin. Polar contacts are indicated by dashed lines in magenta, van der Waals interactions are visualized by showing relevant residues with their (transparent) molecular surface. Superscript capitals designate the G6b-B protein chain. **(C)** LigPlot representation of the heparin-G6b-B contacts, with van der Waals / hydrophobic interactions indicated by the bent comb symbol, and polar contacts shown as dashed lines with distance show indicated in units of Å.

We next measured the binding affinities of G6b-B for the various ligands by surface plasmon resonance. Human G6b-B-Fc-His6 homodimer and human G6b-B-Fc-His6/Fc-StreptagII heterodimer were used as dimeric and monomeric G6b-B molecules, respectively. In the configuration with chip-immobilized G6b-B molecules, heparin bound to both, monomeric and dimeric G6b-B, with high affinity (low nanomolar range). Similar values were obtained for fractionated (9 kDa) HS, and the 12 saccharide heparin oligomer dp12. The binding affinity of perlecan was 366-fold weaker than heparin, in the low micromolar range (Table 3 and Figure 7A). The reverse configuration was also tested, in which ligands were biotinylated and immobilized on streptavidin chips. Binding avidity of dimeric G6b-B to perlecan, fractionated HS and heparin were comparable in this configuration (Table 3 and Figure 7A). Interestingly, a clear difference of monomeric and dimeric G6b-B was observed, with the monomer binding approximately 100-fold weaker than the dimer (Table 3 and Figure 7B) in line with our crystallography data that ligand binding induces dimer formation.

**Table 3.**
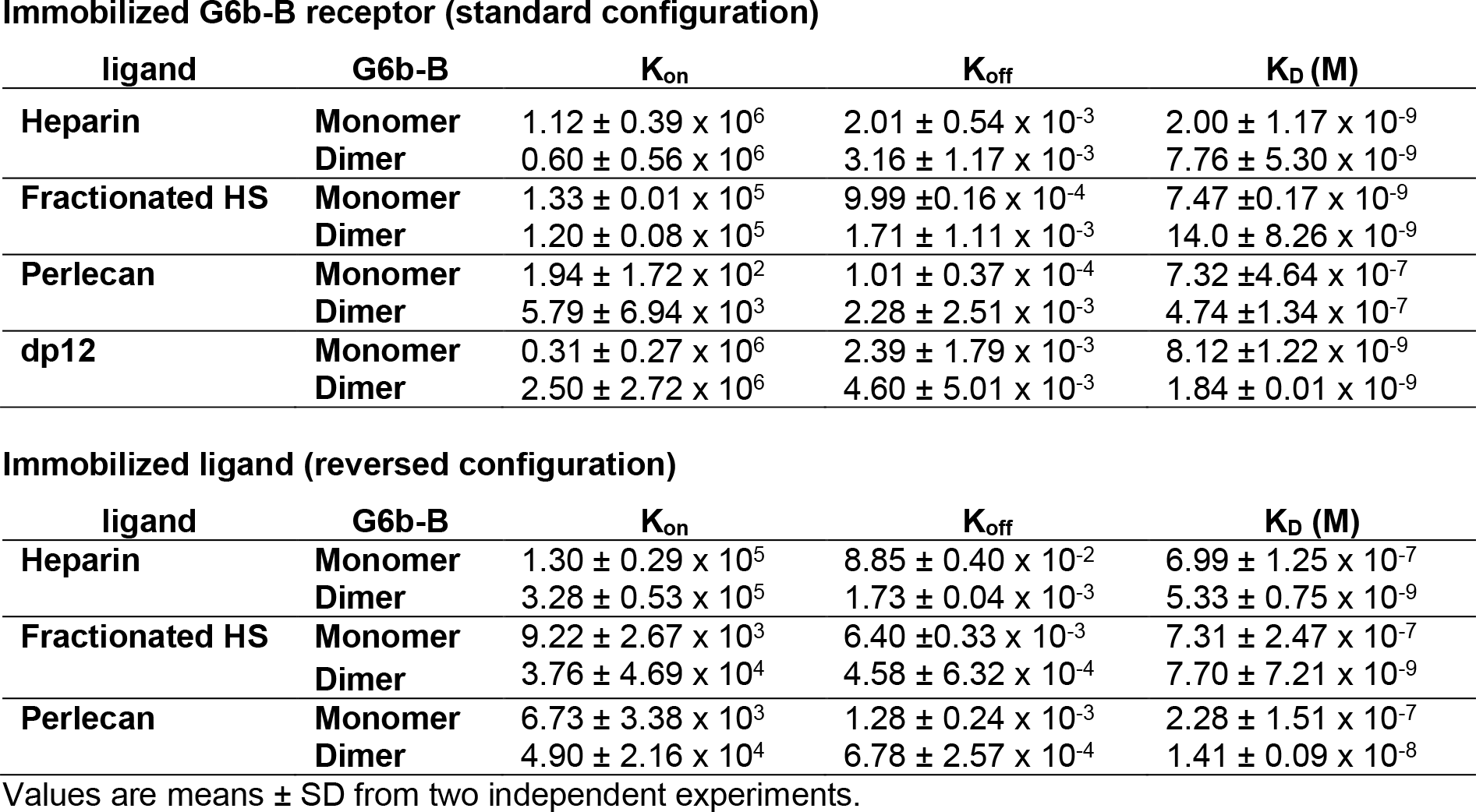
Surface plasmon resonance affinities

**Figure 7.**
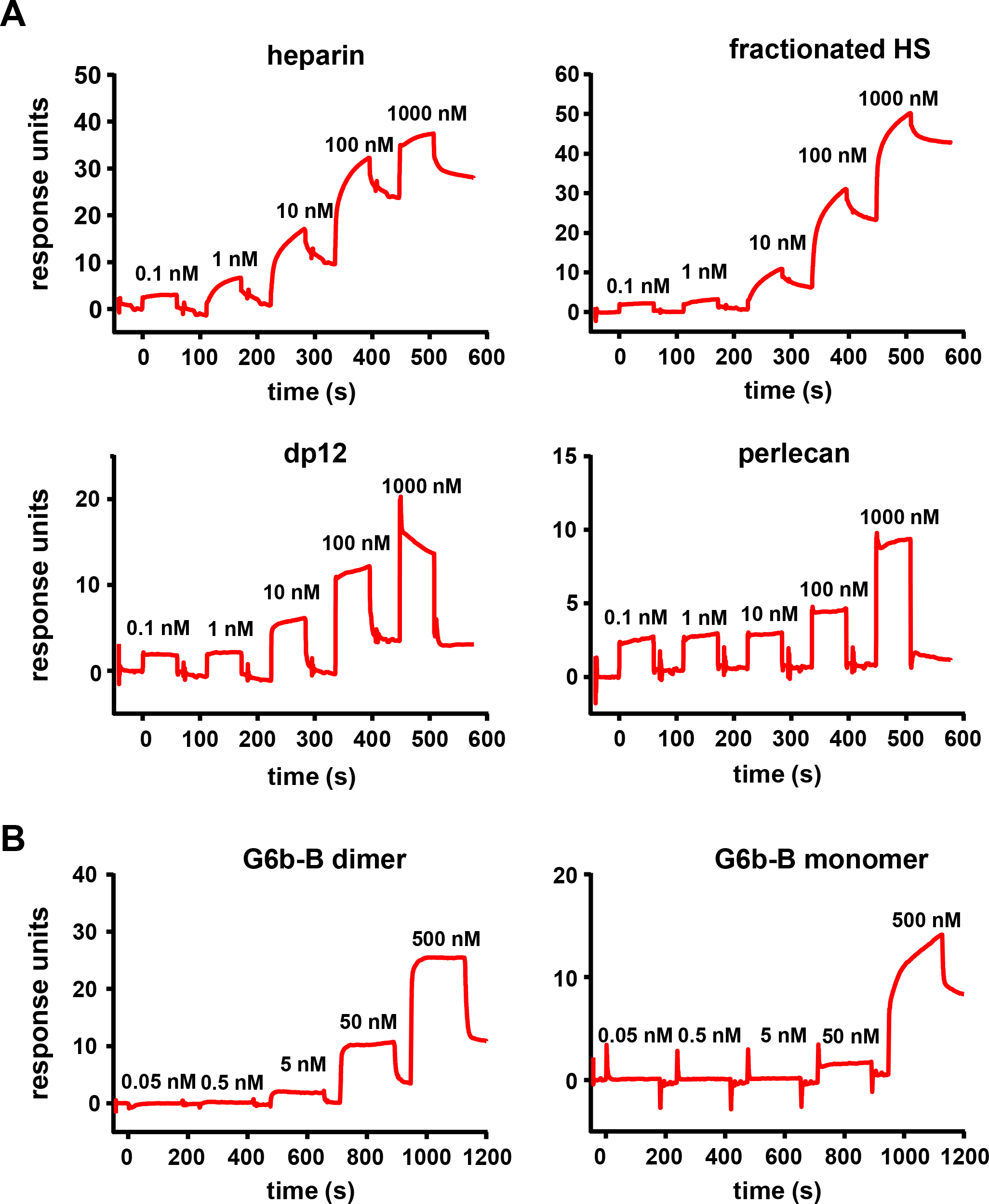
High affinity interaction between G6b-B and its ligands. Representative traces of the surface plasmon resonance experiments, results of which are presented in Table 2. **(A)** Binding of the indicated compound to immobilized dimeric G6b-B in the standard configuration. **(B)** Results from the reversed configuration, depicting traces of dimeric and monomeric G6b-B binding to immobilized heparin.

### Biological effects of perlecan, heparin and HS on platelets and MKs

Having established HS as ligand for G6b-B, we examined the effect of surface bound ligand on platelet function, using an *in vitro* platelet adhesion assay, in which human platelets were incubated on different substrates and their adhesion is quantified colorimetrically. Platelets bound to fibrinogen, as expected, but failed to adhere to perlecan (Figure 8A). However, removal of the HS side-chains by heparinase III treatment resulted in robust adhesion to perlecan. Importantly, perlecan also inhibited the adhesion to fibrinogen and collagen when immobilized together with these substrates. Again, this anti-adhesive effect was abolished upon treatment with heparinase III (Figure 8A). These results suggest that the HS side-chains of perlecan negatively regulate platelet adhesion. To determine whether this inhibitory effect perlecan is mediated via G6b-B, we performed platelet adhesion experiments with platelets from WT and G6b-B knockout (*G6b*^−/−^; KO) mice (Figure 8B). WT mouse platelets exhibited similar adhesion characteristics as human platelets, with the exception that they adhered weakly to heparinase III-treated perlecan (Figure 8B). Platelets from *G6b*^−/−^ mice adhered to fibrinogen similar to WT platelets, however co-coating with perlecan did not inhibit this adhesion, resulting in enhanced adhesion of *G6b*^−/−^ platelets under this condition. Treatment of perlecan with heparinase III abolished this difference (Figure 8B). Adhesion of WT platelets to collagen was inhibited by perlecan in a similar manner as human platelets (data not shown). Platelets from *G6b*^−/−^ could not be meaningfully evaluated on collagen, due to the severe reduction in GPVI surface expression (Mazharian et al., 2012). Collectively, these findings demonstrate that the G6b-B - HS interaction inhibited the adhesion of platelets to the perlecan protein core, collagen and fibrinogen, suggesting an inhibitory on integrin and GPVI signaling.

**Figure 8.**
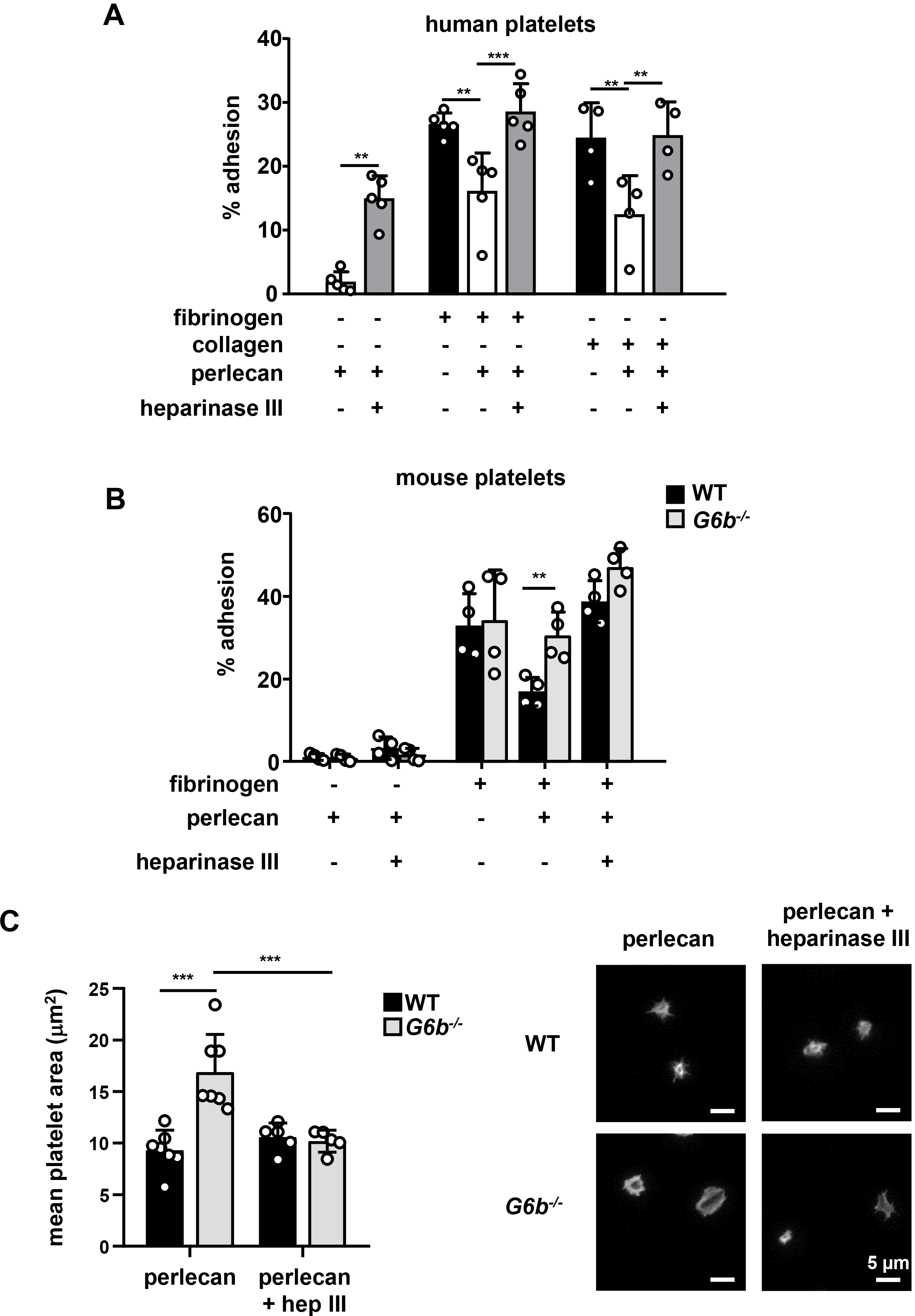
Heparan sulfate removal of perlecan facilitates platelet adhesion. Indicated substrates were coated alone or in combination into 96-well plates (2.5 μg/ml collagen and 10 μg/ml for all other substrates) overnight. Where indicated, wells were treated with 5 mU/ml heparinase III. Platelets from **(A)** human or **(B)** mouse were allowed to adhere for 1 h and adhesion was quantified colorimetrically with pNPP. (A) Human, platelets; n=4-5 individual donors from 3-4 independent experiments; P-values were calculated using one-way ANOVA with Sidak’s post-hoc test; (B) Mouse platelets; n=4 samples/condition/genotype from two independent experiments. Due to severe thrombocytopenia, platelets from up to 5 mice were pooled for one KO sample. P-values for differences between WT and *G6b*^−/−^ mice were calculated using two-way ANOVA with Sidak’s post-hoc test. **(C)** Adhesion of WT and *G6b*^−/−^ platelets on perlecan. (i) Mean surface area of individual platelets quantified by KNIME software analysis, n=5-7 mice/condition/genotype from 2-3 independent experiments using one-way ANOVA with Sidak’s post-hoc test **, P<0.01 and ***, P<0.001. (ii) Representative images of platelets stained for actin with phalloidin-Alexa-488; scale bar: 5 μm; hep III, heparinase III

We next investigated morphological changes in WT and *G6b*^−/−^ platelets adhered to perlecan by microscopy. WT platelets did not spread on perlecan and were small in size (Figure 8C), however, platelets from *G6b*^−/−^ mice spread to a greater extent, indicative for their activation. This effect was abolished upon heparinase III treatment, demonstrating that HS have an activating effect on platelets lacking G6b-B. Of note, platelets from *G6b* diY/F mice, which express physiological levels of a signaling-incompetent G6b-B, recapitulated the enhanced spreading phenotype of *G6b* KO platelets (data not shown). Hence we conclude that G6b-B signaling is required to inhibit platelet activation in the presence of HS.

Staining of mouse bone marrow sections revealed perlecan expression in vessel walls, that co-localized with collagen I and the sinusoid marker endoglin (CD105) (Figure 9A). Thus, MK G6b-B is likely to come into direct contact with HS-side chains of perlecan in sinusoidal vessels during MK maturation and proplatelet formation (Figure 9A). Investigating the impact of the G6b-B interaction with HS on MK spreading, we found that very few WT and *G6b*^−/−^ MKs adhered to a perlecan-coated surface formation (Figure 9B). Whilst perlecan adherent WT MKs were small in size, *G6b*^−/−^ MKs spread to a greater degree on the same substrate. The same effect was observed when perlecan was co-immobilized with fibrinogen and heparinase III treatment abolished the difference (Figure 9B). Hence, similar to platelets, HS resulted in increased cell size in the absence of G6b-B, confirming the inhibitory function of this receptor.

**Figure 9.**
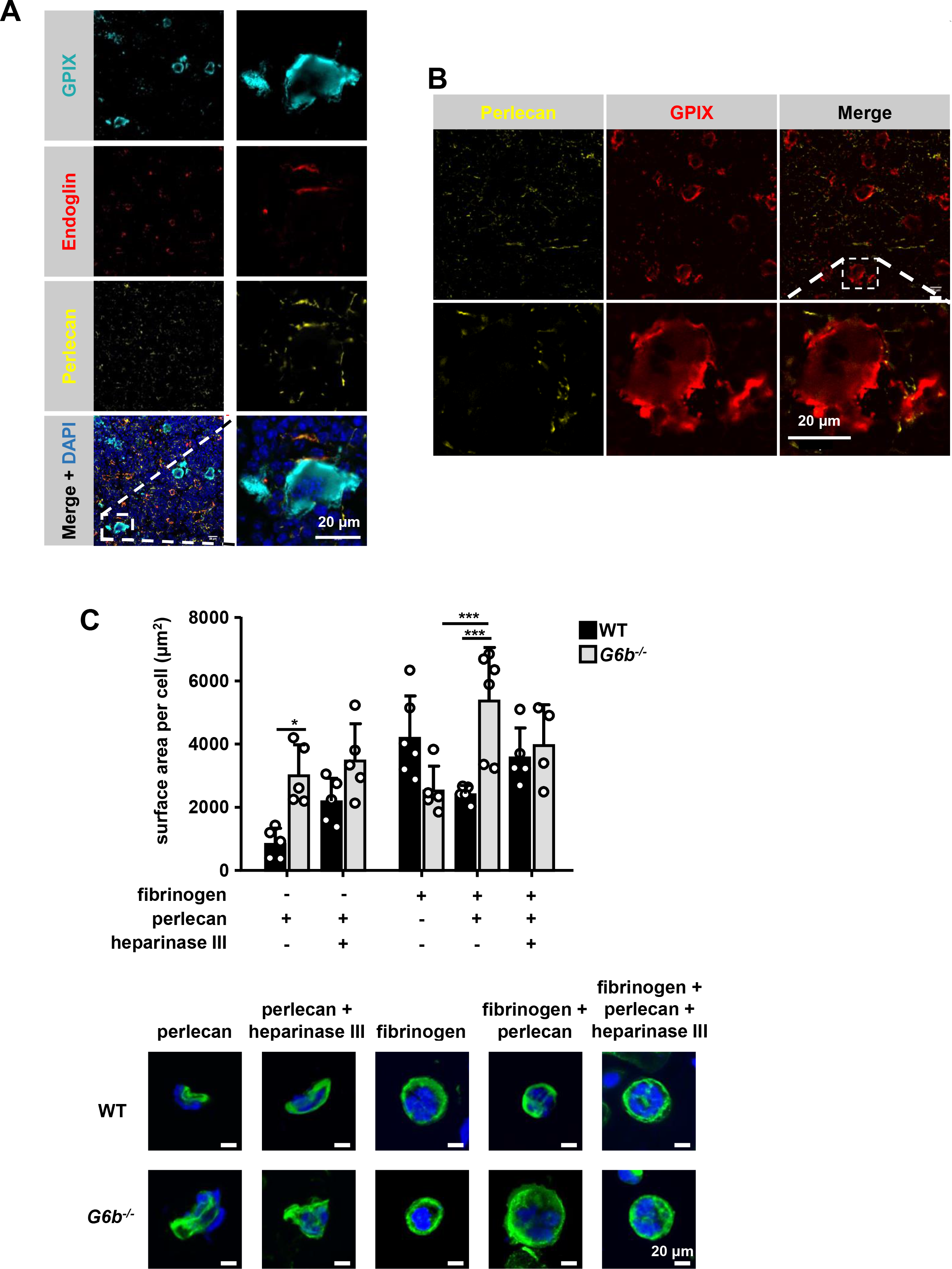
G6b knockout megakaryocytes show enhanced spreading on perlecan. **(A, B)** Immunohistochemical analysis of murine femur sections. Sinusoids were marked using anti-endoglin (CD105) and MKs by anti-GPIX. **(A)** Perlecan is abundantly expressed within the bone marrow cavity and present in intersinusoidal spaces and part of basement membranes in sinusoids and arterioles. MKs (stained with GPIX) come into contact with perlecan. **(B)** Perlecan is not detected inside MKs; scale bar: 20 μm. **(C)** Adhesion of WT and *G6b*^−/−^ MKs on perlecan. (i) Mean surface area of MKs was quantified with ImageJ. n=4-6 mice/condition/genotype from 3 independent experiments; P values were calculated using two-way ANOVA with Sidak’s post-hoc ***, P<0.001; *, P<0.05 (ii) Representative images of platelets stained for tubulin (green) and DAPI (blue); scale bar: 20 μm

We next investigated the biological effects of G6b-B ligands on platelet aggregation in response to collagen, which activates platelets via the ITAM-containing receptor complex GPVI-FcR γ-chain (Nieswandt & Watson, 2003). Heparin and HS both enhanced platelet aggregation in response to subthreshold concentrations of collagen (Figure 10). This is in line with previous reports and may be explained by binding of these ligands to multiple platelet receptors (Gao et al., 2011; Saba, Saba, & Morelli, 1984; Salzman, Rosenberg, Smith, Lindon, & Favreau, 1980 resulting in an overall aggregation-promoting response. We did not find an effect of perlecan on collagen-mediated platelet aggregation at concentrations tested, suggesting that perlecan must be immobilized to surface in order to provide HS chains at a sufficient density to observe the inhibitory effects observed in adhesion experiments (Figure 8,9). In addition, multiple direct and indirect effects on platelets via the perlecan protein core, as described previously (Bix et al., 2007), may mask an effect of the HS chains in this assay.

**Figure 10.**
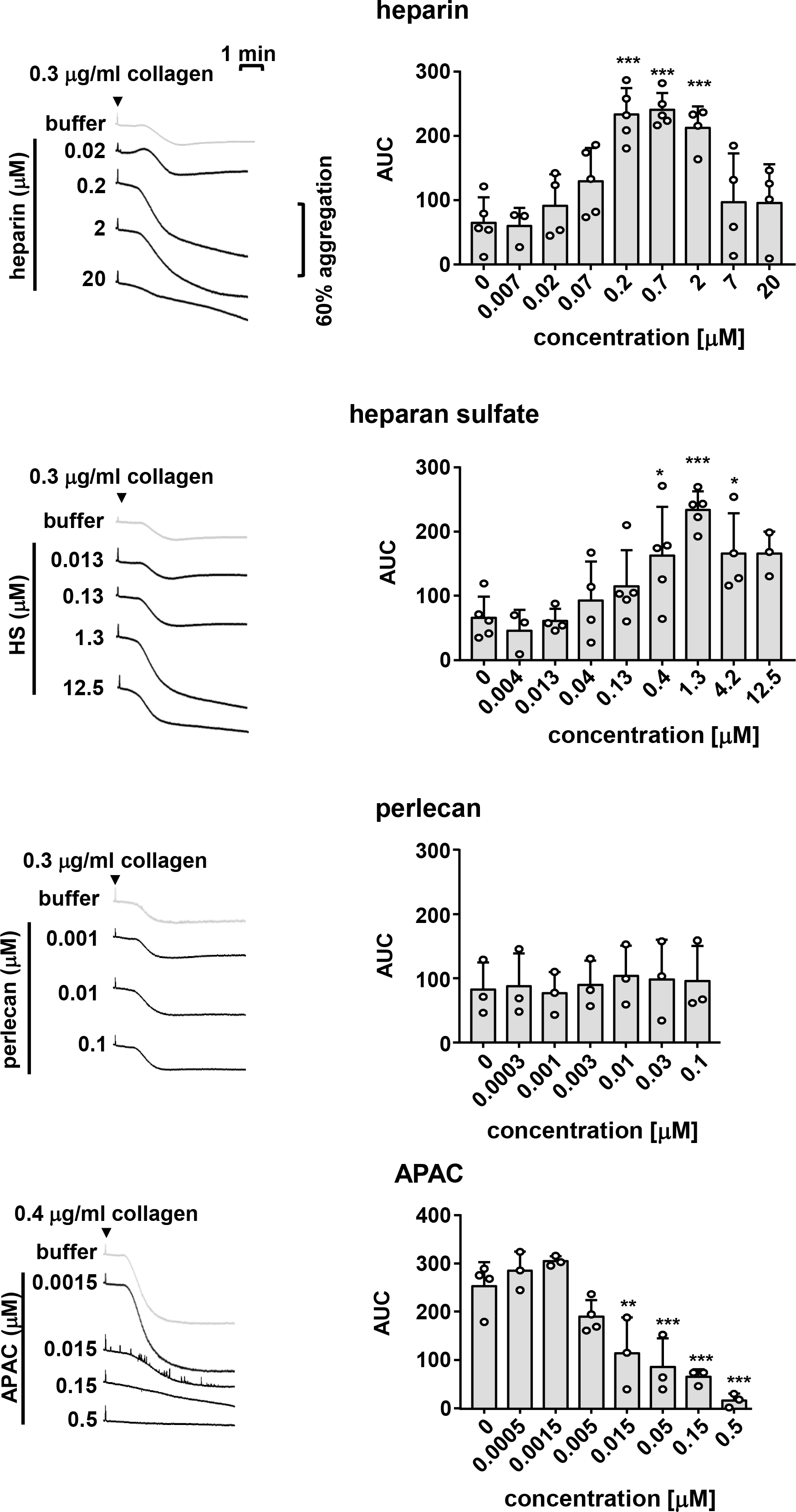
Effects of G6b-B ligands on platelet aggregation. Human platelet rich plasma (PRP) was incubated with the indicated compound for 90 s prior to agonist addition. Aggregation traces were recorded on a Chronolog four channel aggregometer. Averaged aggregation traces (left) and area under the curve (AUC) quantification (right) of platelet aggregation (n=3-5 per condition), P-values were calculated using one-way ANOVA with Dunnett’s post-hoc test and refer to the untreated control; ***, *P*<0.001, **, *P*<0.01 and *, *P*<0.05

To overcome this limitations we took advantage of the multivalent semisynthetic heparin proteoglycan mimetic APAC (Lassila & Jouppila, 2014; Lassila et al., 1997) in this assay. APAC consists of unfractionated heparin covalently coupled to a human albumin core, providing a high local density of heparin molecules. In contrast to single-chain heparin, APAC dose-dependently inhibited collagen-induced platelet aggregation (Figure 10), with an almost complete block observed at 0.5 μM, as previously described (Lassila & Jouppila, 2014).

We next examined the effect of heparin and APAC on WT and G6b-B deficient platelets using a flow cytometric approach, sufficing much smaller sample volumes than aggregation assays, using integrin αIIbβ3 activation (fibrinogen-A488 binding) and degranulation-dependent TLT-1 surface exposure (Smith et al., 2018) as markers for platelet activation. While APAC and heparin had no detectable effect on WT platelets, APAC induced robust integrin activation and platelet secretion, demonstrating a platelet-activating effect of this compound in the absence of G6b-B (Figure 11A and Figure 11-figure supplement 1A). Next we aimed to investigate the impact of G6b-B ligands on ITAM-mediated platelet activation in WT and *G6b* KO mice. Due to severe reduction of GPVI receptor levels in G6b-B deficient animals, resulting in a lack of response to GPVI agonists in this assay (Mazharian et al., 2012), we were not able measure the impact on GPVI-mediated platelet activation. Therefore, we stimulated platelets with an antibody directed against the hemi-ITAM receptor CLEC-2, which is expression is not affected by G6b-B deficiency. APAC, but not heparin, significantly inhibited platelet degranulation in response to CLEC-2 in WT platelets. Importantly, this inhibitory effect of APAC, was not observed in platelets from G6b-B deficient animals (Figure 11B), and was also absent in the platelets from G6b-B diY/F mice, expressing signaling-incompetent form of G6b-B (Figure 11-figure supplement 2). Hence, we conclude that APAC suppresses CLEC-2-mediated platelet activation by inducing an inhibitory signal via G6b-B. Of note, fibrinogen binding to CLEC-2-stimulated platelets was significantly reduced by APAC, in both WT and G6b-B deficient mice, suggesting that APAC might interfere with fibrinogen binding in this experimental setting (Figure 11-figure supplement 1B). Overall, these findings demonstrate that multivalent G6b-B ligands inhibit platelet activation via (hemi)ITAM receptors, while soluble single-chain molecules do not.

**Figure 11.**
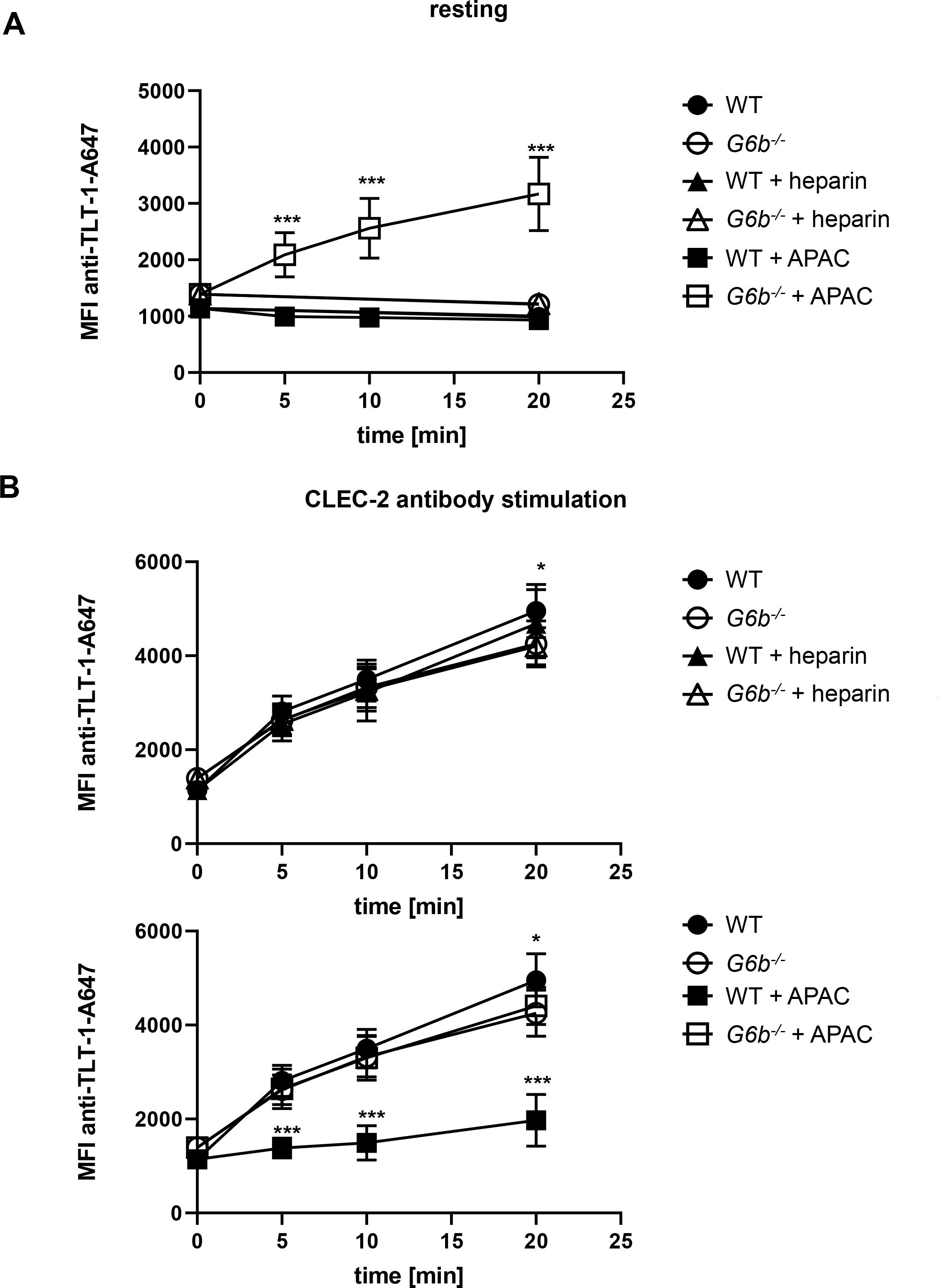
APAC inhibits CLEC-2-mediated degranulation in WT but not *G6b* KO platelets. Mouse blood was incubated with the indicated compounds (0.05 μM) in the **(A)** absence or **(B)** presence or of a stimulating CLEC-2 (3 μg/ml) for the indicated time. Samples were fixed and TLT-1 surface levels, a marker for platelet degranulation were determined by flow cytometry. n=5-6 mice/condition/genotype from 2 independent experiments. P-values were calculated using (A) two-way ANOVA with Sidak’s post-hoc test (comparison of WT APAC vs *G6b*^−/−^ APAC) or (B) two-way ANOVA with Turkey’s post-hoc test and refer to the difference between WT and *G6b*^−/−^ ***, *P*<0.001 and *, *P*<0.05

### Conjugated heparin induces phosphorylation of G6b-B and downstream signaling

We performed signaling studies, to gain mechanistic insights on the opposing effects of soluble heparin vs. conjugated heparin. Washed human platelets were incubated with heparin or APAC, and their lysates were immunoblotted with an anti-phospho-tyrosine antibody (p-Tyr). Both compounds induced moderate changes in whole cell tyrosine phosphorylation as compared to collagen, with APAC having a stronger effect (Figure 12A). The most pronounced change observed in response to G6b-B ligation was an increase in signal intensity of a 150 kDa protein as well as a doublet in the heparin- and APAC-treated sample migrating at 27 and 32 kDa, correlating with glycosylated and non-glycosylated human G6b-B. Hence, we assessed the phosphorylation status of G6b-B using custom phospho-tyrosine-specific G6b-B antibodies, directed against phosphorylated ITIM and ITSM of G6b-B and by immunoprecipitating the receptor and blotting with the p-Tyr antibody (Figure 12A and Figure 12–figure supplement 1). G6b-B was found to be phosphorylated at low levels and have some associated Shp1 and Shp2 in resting platelets, which was enhanced following collagen activation (Figure 12A and Figure 12– figure supplement 1), as previously reported (Mazharian et al., 2012; Senis et al., 2007). Heparin and to a greater extent, APAC, induced phosphorylation of G6b-B, accompanied by an increase in Shp1 and Shp2 association (Figure 12A and Figure 12–figure supplement 1). Similar results were obtained with HS, but to a lesser extent than either heparin or APAC (Figure 12–figure supplement 2). Perlecan did not induce phosphorylation of G6b-B, in line with our observation in the aggregation assay, suggesting that it must be surface-immobilized to have an effect on platelets (Figure 12–figure supplement 1)

**Figure 12.**
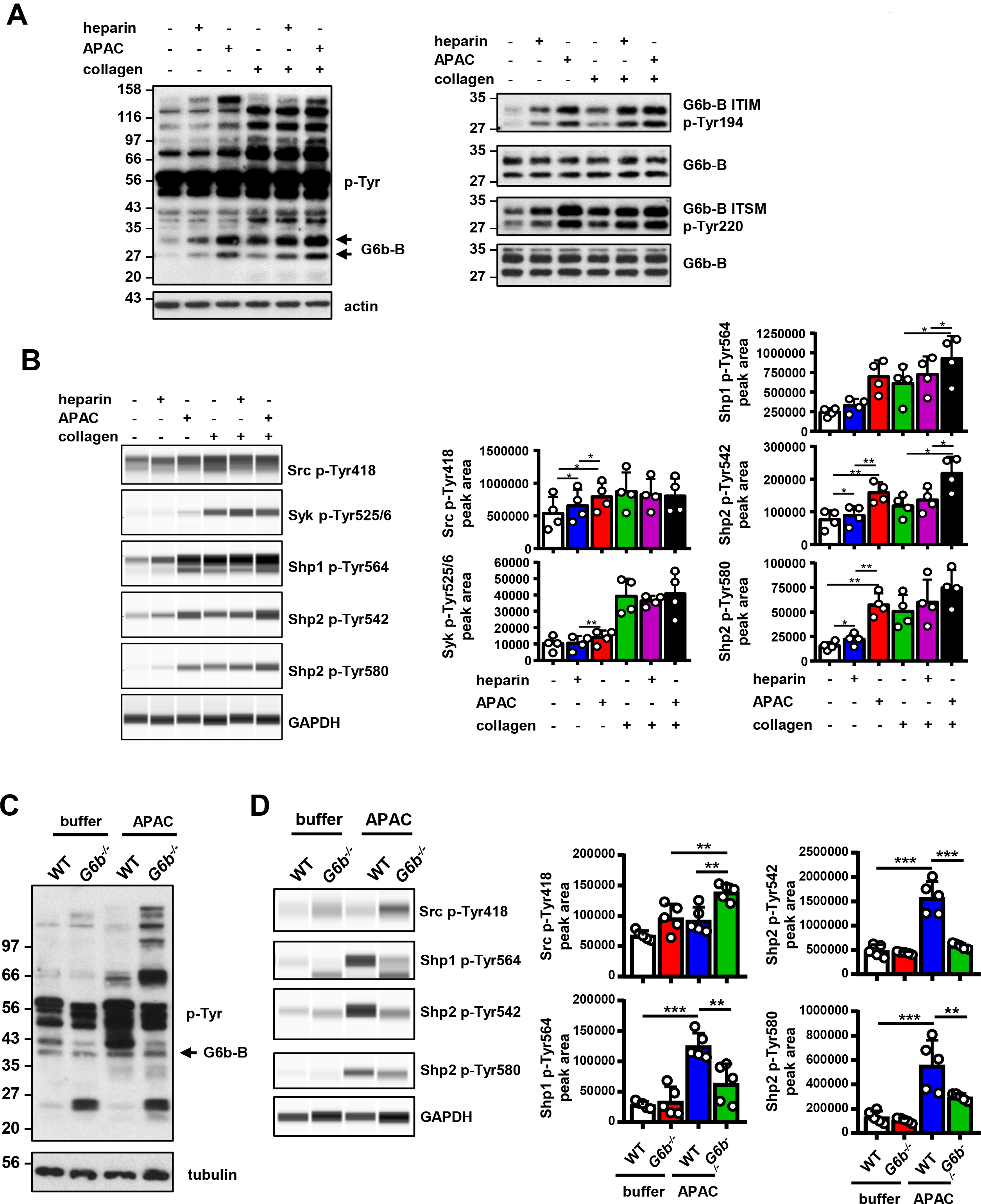
APAC induces G6b-B phosphorylation and downstream signaling. **(A)** Washed human platelets (5 × 10^8^/ml) were incubated for 90 s with 0.05 μM APAC, 0.7 μM heparin or buffer in the presence of 10 μM integrilin. Where indicated, platelets were additionally stimulated with 3 μg/ml collagen for 90 s following compound treatment. Samples were lysed and whole cell lysates (WCL) were analyzed by western blotting. Representative western blots from n=3-5 independent experiments. **(B)** Lysates were also analyzed by quantitative capillary-based gel electrophoresis with the indicated antibodies. Representative data is displayed as blots on the left and quantification of peak areas on the right. **(C, D)** Washed mouse platelets (5 × 10^8^/ml) were incubated for 90 s with 0.05 μM APAC or buffer in the presence of 10 μM lotrafiban. Samples were analyzed as described above. The *G6b*^−/−^ samples show IgG light chain fragments, due to IgG binding to the platelet surface. P-values were calculated using one-way ANOVA with Sidak’s post-hoc test. ***, *P*<0.001, **, *P*<0.01 and *, *P*<0.05; p-Tyr, anti-phosphotyrosine (4G10).

Using a quantitative capillary-based gel electrophoresis platform (ProteinSimple Wes), we investigated the effects of heparin and APAC on the phosphorylation status of the tyrosine phosphatases Shp1 (pTyr562) and Shp2 (p-Tyr580 and p-Tyr542), which are essential effectors of G6b-B signaling (Geer et al., 2018). Strikingly, APAC induced prominent phosphorylation of Shp1 and Shp2, whereas heparin only induced modest changes in Shp2 phosphorylation (Figure 12B). We also observed a marginal increase in SFK phosphorylation (p-Tyr418) in platelets treated with heparin and APAC, correlating with increased phosphorylation of G6b-B under these conditions (Figure 12B).

Subsequently, we compared the effects of heparin and APAC on GPVI signaling in response to an intermediate concentration of collagen (3 μg/ml). Despite both compounds further enhancing collagen-induced phosphorylation of G6b-B, and in case of APAC also phosphorylation of Shp phosphatases (Figure 12B), whole cell phosphorylation remained largely unaltered. Similarly, we also found no inhibitory effect of heparin or APAC on Src (p-Tyr418) and Syk (p-Tyr525/6) phosphorylation, both indirect markers of SFK and Syk activation, and critical kinases for initiating and propagating GPVI signaling (Senis, Mazharian, & Mori, 2014) (Figure 12B).

To corroborate that the APAC-induced increase in Shp1 and Shp2 phosphorylation are mediated by G6b-B, we conducted signaling experiments in platelets from WT and *G6b*^−/−^ mice. APAC treatment of WT platelets recapitulated the effects observed in human platelets, showing only a modest change in overall phosphorylation pattern, and an increase in Shp1 and Shp2 phosphorylation (Figure 12C, D). In contrast, APAC-induced robust tyrosine phosphorylation in G6b-B-deficient platelets (Figure 12C), indicative of reduced inhibitory signaling and platelet hyperreactivity the absence of G6b-B. Strikingly this accompanied by reduced tyrosine phosphorylation of Shp1 and Shp2 in these platelets compared with WT platelets (Figure 12D). Collectively, these findings demonstrate that heparin and APAC have a direct effect on G6b-B phosphorylation, however, only the high-density ligand APAC is able to induce robust downstream inhibitory signaling via G6b-B, culminating in Shp1 and Shp2 binding and tyrosine phosphorylation.

## DISCUSSION

In this study, we present evidence that establishes G6b-B as a functional receptor of HS and heparin. Little was known about the effects of GAGs on platelet and megakaryocyte function and the underlying molecular mechanisms, thus these findings represent a major advance in our understanding of the interaction, biological and biochemical effects of GAGs on these cells. Using a mass-spectrometry-based approach and subsequent *in vitro* binding assays, we identified the HS chains of perlecan as a physiological binding partner of G6b-B. The binding of G6b-B to HS was corroborated by a cell-based CRISPR KO screening, which identified molecules involved in the HS synthesis pathway as a prerequisite of G6b-B binding. Two possible explanations why this assay did not identify perlecan, nor any other individual HSPGs as binding partners of G6b-B: firstly, the CRISPR screening approach will not identify genes that are essential for cell viability, and secondly, it will not identify proteins that have redundant functions. Given that perlecan is secreted from endothelial and smooth muscle cells, it is possible that there could be HSPGs other than perlecan (syndecans / glypicans) on the cell surface that carry the GAG chains in HEK cells. Since the molecules in the HS synthesis pathway are essential for their respective synthesis, they can be identified in this approach more easily. This potential redundancy of HSPGs may also apply for the situation *in vivo*, and we cannot exclude the possibility that G6b-B may interact with other HSPGs in the cardiovascular system.

As with many other HS-binding molecules, G6b-B also binds structurally-related heparin (Xu & Esko, 2014). Indeed, the interaction between heparin and G6b-B had been described previously, but molecular details of the interaction and their functional significance had not been determined (de Vet et al., 2005). Our size exclusion chromatography data demonstrate that dimerization of G6b-B is induced by the heparin ligand. The crystal structure of heparin-bound G6b-B reveals the mode of ligand binding and how binding of this ligand induces ectodomain dimerization. The contact surfaces between the G6b-B dimer and the Fab fragments are spatially separated from the heparin binding site, suggesting that the presence of the Fab fragments does not interfere with heparin binding. Heparin-dependent, non-constitutive dimerization of G6b-B is consistent with the small interface between the G6b-B subunits and the absence of main chain-main chain hydrogen bonds across the β-sheet of the binding surface. Among 34 entries currently in the PDB of structures containing heparin as ligand, dimeric assemblies (or multimeric assemblies with a 2-fold rotation axis) are common (Supplemental Figure 1), but the anti-parallel alignment of 2 Ig-like domains in the heparin-bound structure of G6b-B appears to be unique (Cai et al., 2015; Dahms, Mayer, Roeser, Multhaup, & Than, 2015; Fukuhara, Howitt, Hussain, & Hohenester, 2008; Pellegrini, Burke, von Delft, Mulloy, & Blundell, 2000; Schlessinger et al., 2000). The involvement of the β-sheet surface in heparin binding is somewhat reminiscent of how carbohydrate-binding modules (CBM) bind saccharide ligands (Abbott & van Bueren, 2014). CBMs are non-enzymatic domains often associated with carbohydrate active enzymes, contributing to carbohydrate binding and discrimination (Boraston, Bolam, Gilbert, & Davies, 2004).

The crystal structure of G6b-B shows a prominent positively charged electrostatic surface area, but this positive surface patch runs perpendicular to the central cleft of the G6b-B dimer. Indeed, the heparin oligosaccharide lines up with the cleft, rather than extending along the positive surface patch. Comparison with other heparin-bound structures (Supplemental Figure 1) suggests that charge complementation is not the sole determinant of the mode of heparin binding, but that depth and shape of the docking site are likely to be important as well. Nevertheless, charge complementing ionic interactions lock the ligand into register at the center of the G6b-B binding cleft, whereby the utter sparsity of sulfate-Arg or sulfate-Lys interactions is surprising. The crystal structure rationalizes the diminished binding of G6b-B transfected HEK293 cells to biotinylated heparin when the four basic residues Lys54, Lys58, Arg60 and Arg61 are simultaneously mutated. Among these four side-chains, the key interaction appears to be with Arg60, as Arg61 is shielded through G6b-B dimerization from the ligand, Lys54 is well separated from the binding cleft, whilst Lys58 is situated within a 4 Å-radius of heparin, but makes no polar interactions. The heparin ligand does not exhaust the possibilities for specificity-determining interactions with G6b-B in the ligand-binding cleft. For instance, Arg60^F^, Lys109^F^ are involved in ionic interactions with the same sulfate group, but not their counterparts in chain E on the opposite side of the cleft. It is conceivable that the physiological HS ligand of G6b-B may have a different pattern of sulfate groups that engage both pairs of Arg60, Lys109, perhaps in addition to Lys58.

Investigating the functional consequences of this interaction revealed that heparin and HS have complex effects on platelet function and that G6b-B is a key regulator in this process. Our data demonstrates that, to induce robust inhibitory biological or signaling effects, G6b-B ligands either need to be immobilized to a surface, as in the case of perlecan-coated plates, or multivalent, as in the case of APAC. In contrast, single chain heparin and HS enhanced rather than inhibited platelet aggregation. These findings are in line with numerous previous reports, showing enhancing effects of heparin on platelet aggregation in platelet-rich plasma (Gao et al., 2011; Saba, Saba, & Morelli, 1984; Salzman, Rosenberg, Smith, Lindon, & Favreau, 1980). This most likely also contributes to a mild drop in platelet counts in patients receiving heparin, referred to as non-immune heparin-induced thrombocytopenia (Cooney, 2006). Based on our signaling data and size exclusion chromatography data, we assume that heparin, despite being able to dimerize the receptor, fails to cluster G6b-B sufficiently into higher-order oligomers to induce robust downstream signaling (Figure 13A, B). It remains to be determined, whether the enhancing effects of heparin and HS on platelet aggregation is mediated by inhibiting inhibitory effects of G6b-B alone, or by additional effects on other platelet receptors, which promote platelet activation, such as the integrin αIIbβ3, previously shown to bind heparin (Fitzgerald, Leung, & Phillips, 1985; Gao et al., 2011; Sobel et al., 2001).

**Figure 13.**
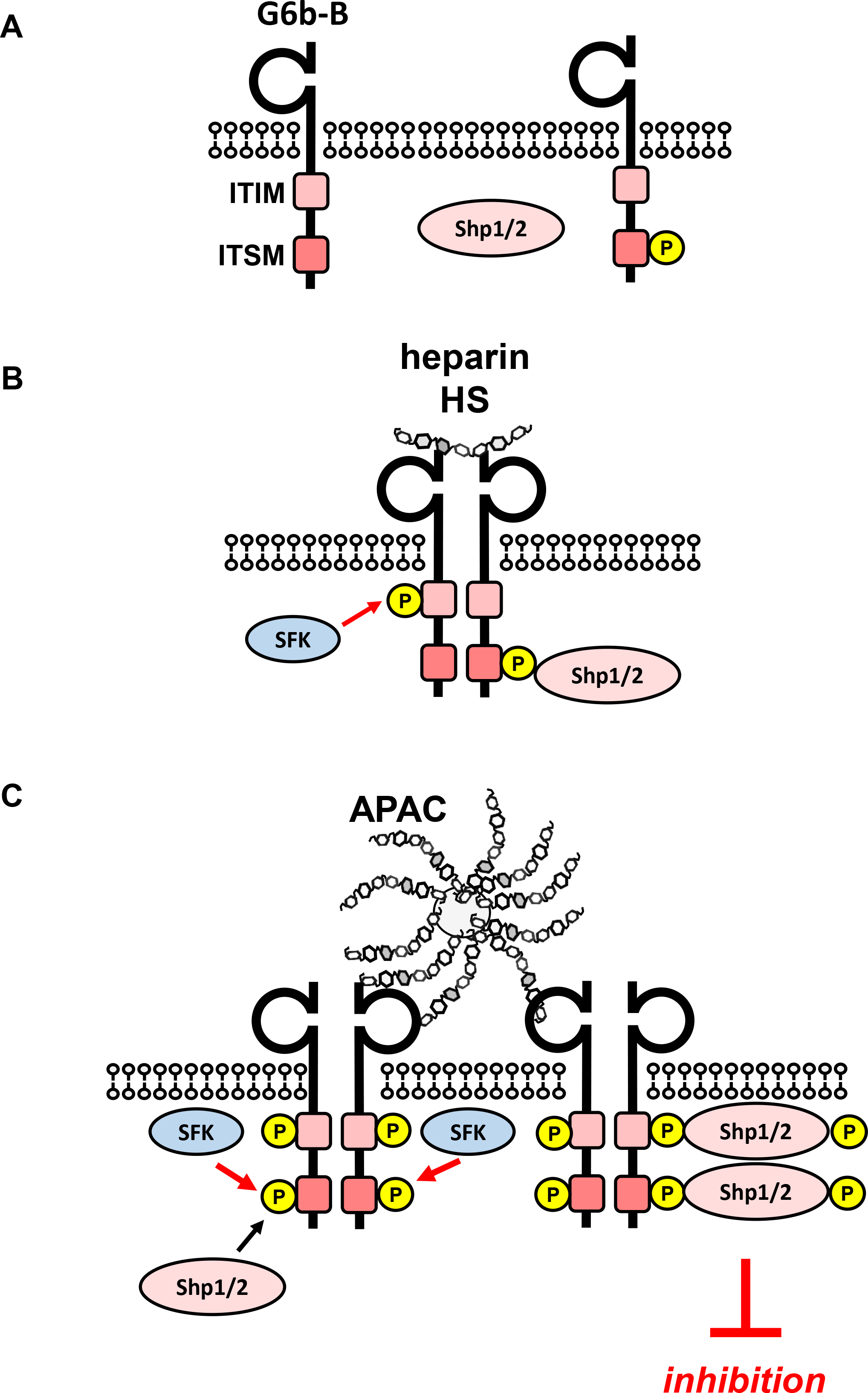
Model of glycan-mediated regulation of G6b-B function. **(A)** In the absence of any ligand, G6b-B is mainly present in a monomeric state and phosphorylated to a low degree. **(B)** Small soluble ligands, e.g. heparin, induce dimerization of the receptor, however, induce only mild G6b-B phosphorylation and downstream signaling. **(C)** Multivalent ligands, e.g. the HS chains of vessel wall perlecan (not shown) or the heparin molecules in APAC, cluster G6b-B dimers into higher order oligomers, hence facilitating downstream signaling of G6b-B, including robust phosphorylation of G6b-B and downstream Shp1 and Shp2 phosphatases, resulting in inhibition of platelet activation. SFK, src family kinase

In contrast to these soluble, monovalent ligands, the HS of immobilized perlecan exerted an inhibitory effect on platelets, as evidenced by impaired adhesion of platelets to collagen and fibrinogen. This extends observations from previous reports describing the anti-adhesive properties of the HS chains of perlecan, although the underlying mechanism was not known (Klein, Conzelmann, Beck, Timpl, & Muller, 1995; Lord et al., 2009). Moreover, heparinized polymers showed less platelet adhesion than their non-heparinized counterparts (Han, Jeong, & Kim, 1989; Lindhout et al., 1995; Olsson, Lagergren, Larsson, & Radegran, 1977). Our results with platelets from G6b-B-deficient mice demonstrate that heparin or HS engagement by G6b-B on these surfaces induce an inhibitory signal, blocking platelet activation and adhesion.

The failure of perlecan in solution to have any effect on collagen-mediated platelet aggregation and platelet signaling, suggests that perlecan must be immobilized to surface in order to provide HS chains at a sufficient density to induce the inhibitory effects observed on platelet adhesion. To determine the effect of G6b-B clustering in solution, we took advantage of APAC, which mimics naturally occurring macromolecular heparin proteoglycans and harboring a higher GAG density than perlecan. Similar to previous reports (Kauhanen, Kovanen, & Lassila, 2000; Lassila & Jouppila, 2014; Lassila et al., 1997), we found that APAC inhibited platelet activation via the ITAM-containing GPVI-FcR γ-chain receptor complex, but also towards the hemi-ITAM-containing receptor CLEC-2. Thus, by increasing the clustering capacity of heparin to a multivalent form, an inhibitory effect on platelet function was achieved in solution. In line with this observation, we found that APAC induced stronger phosphorylation of G6b-B, which was accompanied by association and phosphorylation of the tyrosine phosphatases Shp1 and Shp2, not observed in G6b-B-deficient platelets. We therefore conclude, that clustering of G6b-B receptor dimers into higher-order oligomers by an immobilized or multivalent ligand is required to have an inhibitory effect on platelet function (Figure 13C)

Perlecan is secreted by endothelial and smooth muscle cells into the extracellular space of the vessel wall and hence inaccessible by platelet G6b-B in an intact blood vessel (Murdoch, Liu, Schwarting, Tuan, & Iozzo, 1994; Saku & Furthmayr, 1989; Segev, Nili, & Strauss, 2004). Only upon vascular injury will the interaction between platelet G6b-B and perlecan occur. Given the results of our adhesion assay, we speculate that the interaction of platelet G6b-B with perlecan negatively regulates the initial steps of thrombus formation, preventing thrombi from forming unnecessarily. Our results also demonstrated that platelets and MKs from G6b-B-deficient mice showed an activation response towards the HS chains of immobilized perlecan and, in case of platelets, also towards APAC, even in the absence of a classical platelet agonist such as collagen. Hence, one of the key functions of G6b-B *in vivo* may not solely be restricted to inhibit platelet function upon vascular injury, but also to retain platelets in a resting state. Notably, the HSPGs syndecan-1 and −4 expressed on the surface of endothelial cells that form an integral part of the glycocalyx (Marki, Esko, Pries, & Ley, 2015). As platelets marginate to the vessel wall, the interaction of G6b-B on circulating platelets with the glycocalyx may induce a low level inhibitory signal helping to maintain platelets in an inactive state, in line with basal phosphorylation of G6b-B in resting platelets.

The G6b-B-HS interaction may also be relevant for triggering directional formation of proplatelets by MKs towards sinusoidal blood vessels at sites of platelet production. A key yet unresolved question is how MKs remain relatively refractory and do not release platelets into the ECM-rich environment of the bone marrow despite expressing the same repertoire of cell-surface receptors as platelets. G6b-B is highly expressed in mature MKs and G6b-B KO and loss-of-function mice show a severe macrothrombocytopenia due to impaired proplatelet formation and platelet production (Geer et al., 2018; Mazharian et al., 2012). Here we have demonstrated that G6b-B-deficient, but not WT MKs increase their size in the presence of perlecan, indicative of cellular activation. This is also in line with our observation of large clusters of atypical MKs in the bone marrow of G6b-deficient patients (Hofmann et al., 2018). In addition, our and previous findings show that perlecan is abundantly expressed in the bone marrow ECM and comes into contact with mature MKs ((Farach-Carson, Warren, Harrington, & Carson, 2014), raising the possibility that the MK G6b-B-HS interaction plays a critical role in regulating polarized proplatelet formation into sinusoidal blood vessels.

In summary, our findings establish the interaction of G6b-B with heparan sulfate as a novel mechanism regulating platelet reactivity, as well as having important implications in the regulation of platelet production and the adverse effects observed upon soluble heparin administration.

## EXPERIMENTAL PROCEDURES

### Mice

*G6b* (*G6b*^−/−^) and *G6b diY/F* KI (*G6b*^*diYF/diYF*^) mice were generated on a C57BL/6 background by Taconic Artemis (Cologne, Germany) as previously described (Geer et al., 2018; Mazharian et al., 2012). Control mice were pure C57BL/6 (*G6b*^+/+^), referred to as WT. All procedures were undertaken with the U.K. Home Office approval in accordance with the Animals (Scientific Procedures) Act of 1986.

### Reagents and antibodies

Perlecan (heparan sulfate proteoglycan), biotinylated heparin, p-nitrophenyl phosphate (pNPP), mouse laminin 111, fibronectin, streptavidin, anti-actin (A4700) and anti-tubulin antibody (T6199) and goat anti-human IgG–HRP antibodies were obtained from Sigma-Aldrich, Dorset, UK. Heparinase III was from AMSBiotechnology, Abingdon, UK. Heparan sulfate, fractionated heparan sulfate, heparin, selectively desulfated heparins and heparin oligomers (degree of polymerization -dp) were from Iduron, Alderley Park, UK. The semisynthetic macromolecular conjugate of unfractionated heparin and a human serum albumin, APAC, was from Aplagon Oy, Helsinki, Finland. Purified human IgG-Fc fragment (IgG-Fc) was from Bethyl Laboratories, Montgomery, Texas, USA. Fibrinogen was from Enzyme Research, Swansea, UK recombinant human agrin (N-terminal part; Thr30-Arg1102) and recombinant mouse syndecan-2 from R&D Biotechnologies, Abingdon, UK. Recombinant human laminins were obtained from Biolamina, Sundbyberg, Sweden. Blocking medium (2.5 % horse serum) and 3,3’-diaminobenzidine tetrahydrochloride (DAB) Peroxidase substrate for immunohistochemistry were purchased from Vector Laboratories, Peterborough, UK and 3,3,5,5-tetramethylbenzidine (TMB) was from BD Biosciences, Wokingham, UK. Polyclonal anti-Shp1 and anti-Shp2 antibodies were from Santa Cruz Biotechnology, Heidelberg, Germany. Polyclonal phospho-specific G6b-B antibodies were generated by Biogenes, Berlin, Germany. Rabbit monoclonal anti-Shp1 p-Tyr564 (D11G5), anti-GAPDH (14C10) and rabbit polyclonal anti-Shp2 p-Tyr542, anti-Shp2 p-Tyr580, anti-Syk p-Tyr525/6 antibodies were from Cell Signaling Technology, Leiden, The Netherlands. Rabbit polyclonal anti-Src p-Tyr418 antibody and phalloidin-Alexa 488 was from Invitrogen Life Technologies, Paisley, UK and anti-phophotyrosine (4G10) from Millipore, Watford, UK. All other antibodies and chemicals were either purchased or generated as previously described (Mazharian et al., 2012).

### Constructs

#### Recombinant proteins

The cDNA encoding the mouse G6b-B extracellular domain was amplified by PCR using the primers GATC AAGCTT ATG GCC TTG GTC CTG CCG CTG (forward) and GATC GGATCC ACT TAC CTG T CTC GTA CCC GTG GGT AGA TCC (reverse) from a mouse megakaryocyte cDNA library template. The PCR product was cleaved using Hind III and Bam HI and ligated into pCDNA3Ig, comprised of the genomic human IgG1 hinge-C2-C3 Fc region cloned into the HindIII and Not I sites of pcDNA3. This creates a construct encoding the extracellular part of G6b, spliced in frame with the IgG1 hinge, producing a G6b-B-Fc chimeric dimer. The resulting protein, mG6b-B-Fc, was expressed in COS-7 cells and then purified via affinity chromatography. The human G6b-B-Fc dimer (hG6b-B-Fc) construct was produced using an identical approach to the murine construct, using the primers GATC AAGCTT ATG GCT GTG TTT CTG CAG CTG (forward) and GATC GGATCC ACTTACCTGT CTG GGG ATA CAC GGA CCC ATG (reverse). Similarly, untagged monomeric G6b-B (residues 18-142) as well as His-tagged versions were produced – human G6b-B (residues 18-142)-Fc-His6 (expressed as a homodimer) and human G6b-B (residues 18-142)-Fc-His6/Fc-streptagII (heterodimer, monomeric for G6b-B; Peak Proteins Limited, Alderley Park) for use in surface plasmon resonance measurements. All human constructs were expressed transiently in HEK293-6E cells.

#### Cell culture

The cDNA encoding the full length of human G6b-B protein was amplified by PCR from a human cDNA library. This PCR fragment was first cloned into the pCR®-Blunt vector (Invitrogen), and then subcloned into the pCDNA3 vector, for expression of untagged G6b-B in heterologous cell systems. Subsequently, the G6b-B mutant with mutation in the potential heparin binding site (hG6b-B K54D/K58D/R60E/R61E referred to as hG6b-B-mut) was generated with the Quick Change Site-directed mutagenesis kit (Agilent Technologies, Stockport, UK).

### Immunohistochemistry

Immunohistochemistry stainings were performed according to standard protocols. In brief, frozen mouse tissue sections (Zyagen, San Diego, CA, UK) were thawed and washed once in phosphate buffered saline (PBS). After blocking for 20 minutes (min) at room temperature (RT), tissues were incubated with mG6b-B-Fc or human IgG-Fc fragment (negative control, 5 μg/ml in PBS) for 75 min at RT. After three washing steps in PBS, slides were fixed in acetone/PFA for 4 min and endogenous peroxidase was blocked with 3% H_2_O_2_ in methanol (5 min). Slides were incubated with anti-human IgG–HRP antibody (1:600 in PBS, 0.1% Tween 20) and the signal developed with DAB substrate. Subsequently, tissue sections were counterstained with hematoxylin. Images were acquired on a Zeiss Axio Scan.Z1 (Zeiss, Cambridge, UK) equipped with an 3CCD color 2MP Hitachi 1200×1600 HV-F202SCL camera, using a 10x (NA 0.45) or 20x (NA 0.8) plan apochromat air objective. Images were acquired and exported with the Zeiss Zen software.

### Femur sectioning and staining

Femurs of mice aged 6–12 weeks were sectioned and stained as described previously (Kawamoto, 2003; Semeniak et al., 2016). In brief, megakaryocytes were stained with anti-GPIX antibody (emfret analytics, Eibelstadt, Germany), CD105 (eBioscicnce) was used as an endothelial cell marker. Additional stainings were performed using antibodies against perlecan (Santa Cruz). Corresponding secondary antibodies detecting IgG of rat, goat, rabbit or mouse were purchased as conjugates with Alexa Fluor 488 (A-11034), 594 (A-11007) or 647 (A-21247, A-21469, A-21244), respectively, from Life Technologies, Darmstadt, Germany, and used at a 1:300 dilution. Slides were mounted in DAPI-containing medium (Southern Biotech, Birmingham, AL, USA). Recording was performed at a Leica TCS SP8 confocal laser scanning microscope (Leica, Wetzlar, Germany) with an 40x oil objective at 20°C. Numerical aperture (NA) of the objective lense was 1.3 and the software used for data acquisition was LASX. Subsequently images were processed with ImageJ (NIH, Bethesda, MD, USA). No 3D reconstruction, gamma adjustments or deconvolution were performed.

### Pull-down and identification of the ligand

Venae cavae were dissected from wild-type mice and fat and connective tissue were removed. The endothelial tissue was placed in lysis buffer (10 mM Tris-HCl, pH 7.6, 150 mM NaCl, 1 mM EGTA, 1 mM EDTA, 1% IGEPALCA-630, 5 mM Na_3_VO_4_, 0.5 mM 4-(2-aminoethyl) benzenesulfonyl fluoride hydrochloride, 5 μg/ml leupeptin, 5 μg/ml aprotinin, 0.5 μg/ml pepstatin) and homogenized with a PowerGen homogenizer (Fisher Scientific, Loughborough). Lysates were centrifuged at 13,000 ×g for 10 minutes at 4°C. Supernatants were collected and re-centrifuged under the same conditions. Protein lysate was precleared with Protein G sepharose (PGS, 50% slurry) and human IgG-Fc fragment by agitation for 1 h at 4°C. The lysate was then split into two samples which received either mG6b-B-Fc or human IgG-Fc fragment (negative control). After 1.5 h PGS was added and samples were agitated for another 1.5 h at 4°C. Finally, PGS was washed three times in lysis buffer and bound proteins were eluted by boiling the PGS pellet for 5 min in 40 μl 2x SDS sample buffer. Samples were then resolved on a NuPage 4-12% Bis-Tris-Gradient Gel (Invitrogen), alongside with mG6b-B-Fc (additional negative control) and stained with colloidal Coomassie. Bands appearing in the mG6b-B-Fc pulldown, but not in the negative controls, were excised and subjected to mass spectrometry analysis (Orbitrap, Thermo Fisher Scientific, Paisley, UK). Corresponding areas from the control pulldown were cut and analyzed in parallel to account for background signals.

### *In vitro* binding assay

Nunc MaxiSorp™ plates (Thermo Fisher Scientific) were coated overnight with 50 μl of substrates, diluted in PBS (supplemented with 0.9 mM CaCl_2_ and 0.5 mM MgCl_2_ for laminins) at a concentration of 5 μg/ml. Plates were washed 3 times with Tris buffered saline (TBS) containing 0.1% Tween 20 (TBS-T) and blocked for 1.5 h at 37°C with 2% fat free milk in TBS, 0.02% Tween 20. For heparin immobilization, biotinylated heparin (5 μg/ml) was added to streptavidin-coated wells for 1 h at RT prior to the blocking step. After one washing step, recombinant G6b-B-Fc or human IgG-Fc fragment (negative control) in 3% BSA in TBS-T was added and incubated for 2 h at 37°C. In competition assays, this incubation step was performed in the presence of the indicated compound. After five washing steps, wells were incubated with HRP-conjugated anti-human IgG antibody for 1 h at RT at low agitation. Alternatively, monomeric, untagged G6b-B was incubated with anti-G6b-B antibody and bound complexes were detected with HRP-conjugated anti-mouse IgG antibody. Plates were washed seven times and signals were developed with TMB. The reaction was stopped by addition of 2 M H_2_SO_4_ (50 μl/well) and absorbance at 450 nm and 570 nm (background) was measured with a Versa max plate reader (Molecular Devices, Wokingham, UK).

### Genome-wide cell-based genetic screening

The cell-based genome-wide genetic screen was performed essentially as described (Sharma et al., 2018). In brief, 3×10^7^ Cas9-expressing HEK293 cells were transduced with a library of lentiviruses each encoding a single gRNA from a pool of 90,709 individual gRNAs targeting 18,009 human genes at a low multiplicity of infection of 0.3 to increase the chances that each cell received a single gRNA. Ten million lentivirally transduced cells were selected using a blue fluorescent protein (BFP) marker three days after transduction using fluorescence-activated cell sorting. The sorted cells were placed back into culture and further selected for five days with 2 μg/mL puromycin. On day nine post transduction, 100×10^6^ cells were stained with a recombinant protein consisting of the entire ectodomain of biotinylated human G6b-B clustered around phycoerythrin (PE)-conjugated streptavidin for an hour at room temperature. The cells were sorted using an XDP flow sorter and the BFP^+^/PE^−^ population collected, representing ~1% of the total cell population. A total of 600,000 cells were collected from which genomic DNA was extracted, gRNA sequences amplified by PCR and their abundances determined by next generation sequencing. The enrichment of gRNA sequences targeting specific genes in the sorted versus unsorted populations were quantified from the sequence data using the MAGeCK software (Li et al., 2014) as previously described (Sharma et al., 2018).

### Surface Plasmon resonance

The interaction of the recombinant heterodimeric (‘monomer’) and homodimeric (‘dimer’) human G6b-B extracellular domain with different ligands was quantified using a BIAcoreTM 8K instrument (GE Healthcare, Little Chalfont, UK). Recombinant G6b-B proteins were immobilized on CM5 sensor chips (GE Healthcare) via an Fc antibody using the Human Antibody Capture Kit (GE Healthcare). Immobilization levels ranged from 7,800-9,000 response units (RU) for the Fc antibody and 3000 to 4000 RU for the G6b-B proteins. Single cycle kinetics (SCK) measurements were undertaken with perlecan, heparin, fractionated HS and dp12. The analytes were injected in increasing concentrations of 0.1, 1, 10, 100 and 1,000 nM. Analytes were flowed over the immobilized G6b-B surface at 30 μl/min with 60 s injection time and 60 s dissociation per concentration. In the ‘reversed configuration’, biotinylated heparin, HS and perlecan were immobilized on streptavidin sensor chips (GE Healthcare); fractionated HS and perlecan were biotinylated using the Lightning-Link^®^ Biotinylation kit (Innova Biosciences, Cambridge, UK). Immobilization levels of the biotinylated species were between 900 and 1000 RU. SCK of ‘monomeric’ and ‘dimeric’ G6b-B were evaluated at 0.05, 0.5, 5, 50 and 500 nM. The analytes were flowed over the immobilized peptides at 10 μl/min with 180 s injection time and 360 s dissociation at each concentration. Data were collected from two replicates per experiment type and analyzed using the BIAevaluation software (GE Healthcare). Sensorgrams were double referenced prior to global fitting the SCK sensorgrams to one-to-one binding models for determining the rate constant of association (k_on_) and dissociation (k_off_). Binding affinities (K_D_) were calculated from the equation KD = k_off/_k_on_.

### Theoretical modeling of G6b-B structure

The G6b-B ectodomain model was generated by submitting the amino acid sequence for G6b-B residues 18-142 to the RaptorX Structure Prediction server (http://raptorx.uchicago.edu/) (Kallberg et al., 2012). Subsequent modelling of the K54D, K58D, R60E, R61E G6b-B mutant and all molecular graphics figure generation was carried out using PyMOL (The PyMOL Molecular Graphics System, Version 2.0 Schrödinger, LLC.). The electrostatic surfaces of both wild-type and mutant G6b-B models were calculated using the APBS suite (Jurrus et al., 2018).

### Crystallography

#### Production of recombinant G6b-B and anti-G6b-B Fab fragment

The G6b-B extracellular domain (ECD) construct encompassing residues 18-133 including the mutations N32D, S67A, S68A, S69A, T71A was expressed in mammalian HEK293 cells and purified by cation exchange and size exclusion chromatography. The recombinant anti-G6b-B Fab fragment was also produced in HEK cells, synthetic genes for light and heavy chains were obtained from Invitrogen GeneArt. The G6b-B ECD-Fab complex was formed by incubating the components together for 2 hours at room temperature with G6b-B ECD in a 1.5 molar excess, and the complex was subsequently purified by size exclusion chromatography. Protein was concentrated to 12 mg/ml in 20 mM Hepes pH 7.1, 75 mM NaCl and finally incubated with 2 mM (10-fold molar excess) of the heparin oligosaccharide dp12 for 1 hour at 4°C prior to setting up the crystallization experiment.

#### Production of crystals and solving of structure

Crystals were grown by vapor diffusion at 20°C in 50 mM MES pH6.2, 10% PEG 550MME, 5% v/v glycerol, and 50 mM CaCl_2_ and appeared within 3 days. Crystals were harvested straight out of the growth drop and cryo-cooled in liquid nitrogen. X-ray diffraction data were collected at 100K on beamline I03 at Diamond Light Source and processed using XDS (Kabsch, 2010) and Aimless (Evans & Murshudov, 2013) via AutoPROC (Vonrhein et al., 2011). The crystal were in space group C2 with the cell dimensions of a=183.80 Å, b=72.34 Å, c=131.04 Å, β=124.52°, and extended to 3.1 Å resolution (Table 3).

The structure was initially solved by molecular replacement using the program Phaser (McCoy et al., 2007) and with a model of the Fab fragment generated from the PDB structure 4K2U (Chen, Paing, Salinas, Sim, & Tolia, 2013) as the search model. This resulted in the placement of two Fab molecules in the asymmetric unit (Phaser Z-score after translation search=10.2). Examination of the resulting electron density maps showed substantial unmodeled density in the vicinity of the CDR regions of both Fab molecules, which was interpreted as bound G6b-B ECD. Multiple rounds of model building in Coot (Emsley & Cowtan, 2004) and refinement using Refmac5 (Murshudov, Vagin, & Dodson, 1997) resulted in a model encompassing about 90% [101 out of 116 residues] of the of G6b-B ECD chain. Residual density at that stage was identified as a single molecule of heparin dp12 bound, with the density covering 8 of the 12 saccharide units in dp12.

The final model represents a complex of G6b-B ECD, dp12 and Fab fragment chains in the ratio 2:1:2 respectively. The refined structure of G6b-B ECD chain has observable electron density for residues Pro19 to Thr38, Arg43 to Arg83 and Ile91 to Cys129. The G6b-B ECD as expected is shown to be a member of the IgV superfamily with the solved structure comprises two antiparallel β-sheets formed by strands ABDE and A′CC′FG. There is also evidence from the electron density for O-linked glycosylation at Thr73 in both copies of the G6b-B ECD. Final refinement statistics for the G6b-B ECD-dp12-Fab dimer complex are given in Table 3.

### Size chromatography of G6b-B ECD

The G6b-B ECD protein encompassing residues 18-133 (N32D, S67A, S68A, S69A, T71A) was either analyzed immediately, or after incubation with heparin oligosaccharide dp12. A Superdex 75 10/300 GL column (GE Healthcare) was both equilibrated and run in 20 mM Hepes, pH 7.1, 75 mM NaCl. dp12 was added to the G6b-B ECD at a 4-fold molar excess (150 μM final concentration). After the addition of dp12 the sample was aspirated gently and incubated for 90 min on ice, prior to SEC analysis. Columns were run at 0.3 ml/min, and 400 μl of G6b-B ECD samples loaded (200 μg). A calibration curve was prepared in the same buffer using conalbumin (75 kDa), ovalbumin (44 kDa), carbonic anhydrase (29 kDa), ribonuclease A (13.7 kDa) and aprotinin (6.6 kDa) (LMM gel filtration standard kit, GE Healthcare). This calibration curve was then used to estimate the molecular weight of both G6b-B ECD and G6b-B ECD +dp12 in order to determine their polymeric states.

### Flow-cytometric analysis of heparin binding transfected CHO cells

Transfections of WT or mutant hG6b-B into CHO cells were carried out in 6-well plates (3×10^5^ cells in 2 ml DMEM medium, supplemented with 10% fetal bovine serum, 2 mM glutamin) using polyethylenimine (Sigma-Aldrich) as described (Ehrhardt et al., 2006). Cells were harvested 2 days after transfection, by detaching them with accutase, and resuspended in PBS containing 0.2 mg/ml BSA and 0.02% sodium azide. Cells were incubated with heparin-biotin (10 μg/ml) and mouse anti-human G6b-B antibody for 45 min at RT, washed twice, and incubated with streptavidin-PE (BD Biosciences) and anti-mouse-alexa488 antibody (Invitrogen). Cells were fixed with 1% formaldehyde and analyzed on a BD FACSCalibur (BD Biosciences).

### Aggregometry

Platelet rich plasma (PRP) was prepared from blood collected from healthy drug-free volunteers as described previously (Dawood, Wilde, & Watson, 2007). Donors gave full informed consent according to the Helsinki declaration. In brief, 9 volumes of blood were collected into 1 volume of 4% (w/v) sodium citrate solution. Blood was centrifuged at 200 ×g for 20 min at RT and PRP was collected. Platelet aggregation was measured using a lumi-aggregometer (Chrono-Log, Abbington On Thames, UK, Model 700).

### Platelet adhesion assay

This assay was performed as described previously (Bellavite et al., 1994). In brief, Nunc MaxiSorp™ plates were coated overnight with 50 μl of substrates, diluted in PBS at a concentration of 10 μg/ml, except for collagen which was used at 2.5 μg/ml. Plates were then washed 3 times and blocked with 2% BSA in PBS for 1 h at 37°C. After washing, 50 μl heparinase III (5 mU/ml) or buffer (20 mM Tris-HCl, pH 7.5, 0.1 mg/ml BSA and 4 mM CaCl_2_) were added to each well and incubated for 1.5 h. After three washing steps, 50 μl of platelet suspension modified Tyrode’s buffer, prepared as previously described (Pearce et al., 2004), at a concentration of 1×10^8^/ml were added and incubated for 1 h at 37°C. After three washing steps with PBS, 140 μl of substrate solution was added to each well and incubated on a rocker at RT for 40 minutes. Then 50 μl of 3M NaOH was added and the signal was quantified 5 minutes later by measuring the absorbance at 405 nm and 620 nm (background). Percentage of adhesion was calculated by normalizing the measured ODs to the signal obtained by directly lysing 50 μl of platelet suspension.

### Flow cytometric analysis of platelet activation

5 μl staining solution, containing 1.5 μg fibrinogen-Alexa488 conjugate (Invitrogen) and 1 μg of anti-TLT-1-Alexa 647 antibody (Biotechne, Abingdon, UK) and 5 μl of whole blood were provided in a well of 96-well plate. Stimulation was started by adding 40 μl of heparin, APAC (0.05 μM final concentration) or buffer, with or without CLEC-2 antibody (3 μg/ml final concentration; Bio-Rad, Oxford, UK). The plate was incubated in the dark for the indicated time and the reaction was stopped by addition of 200 μl 1% ice-cold formalin. Samples were analyzed on a BD Accuri flow cytometer. Platelets were gated using forward and side scatter.

### Preparation and culture of mouse megakaryocytes

Mature megakaryocytes from mouse bone marrow were defined as the population of cells generated with the methodology previously described (Dumon, Heath, Tomlinson, Gottgens, & Frampton, 2006; Mazharian, Ghevaert, Zhang, Massberg, & Watson, 2011).

### Microscopical analysis of platelet and MK adhesion

Glass Coverslips (5 mm diameter) were incubated with 50 μl of perlecan (25 μg/ml), fibrinogen (25 μg/ml) or both overnight at 4 °C. Surfaces were then blocked with denatured BSA (5 mg/ml) for 1 h at room temperature. After washing, 50 μl heparinase III (12.5 mU/ml) or buffer were added to each well and incubated for 1.5 h at 37 °C. Platelets (2×10^7^/ml, 50 μl) were transferred to the slides and incubated at 37 °C for 45 min in a humid atmosphere. Mature megakaryocytes (6×10^3^/ml, 100 μl), were incubated for 5 h. Non-adherent cells were removed by gently washing wells with PBS and adherent cells were fixed with 3.7% paraformaldehyde and permeabilized with 0.2% Triton-X 100 in water. MKs were stained with tubulin-antibody for 1 h followed by anti-mouse-Alexa-488 and rhodamin-conjugated phalloidin for 30 minutes and coverslips were mounted onto microscope slides for imaging using or Antifade Mountant with DAPI (Invitrogen). Platelets were stained with phalloidin-Alexa-488 for 1 h and coverslips were mounted using Hydromount (National Diagnostics, Nottingham, UK). Images were captured by a Zeiss Axio Observer.Z1 / 7 epifluorescence microscope using ZEN Software and 20x (MK) or 63x oil immersion (platelet) plan apochromat objectives.

For platelets, each coverslip was imaged in three random areas. For analysis, the central quarter of each field of view was cropped (1024 × 1024 pixels) and an ilastik pixel classifier software was used to outline a binary segmentation (Sommer, Straehle, Kothe, & Hamprecht, 2011). To distinguish touching platelets, KNIME analytic platform was used to manually identify the centre of individual platelets (Berthold et al., 2009). These coordinates were used to produce the final segmentation of individual platelets, and platelet size was subsequently calculated.

For MK, three tiles of 3×3 images were acquired per coverslip. Average surface area per cell was calculated by analysing total surface area and number of cells per image using ImageJ. Both, imaging and analysis, were done blinded.

### Western Blotting and immunoprecipitation

Washed human or mouse platelets (5×10^8^/ml) or in the presence of 10 μM integrilin or lotrafiban, (integrin αIIbβ3 inhibitors) respectively, were incubated with the respective compound under stirring conditions (1200 rpm, 37°C) for the indicated time. Platelets were lysed by addition an equal volume of ice cold 2 × lysis buffer an insoluble cell debris was removed by centrifugation for 10 minutes at 13,000 × g, 4°C.

For immunoprecipitations, whole cell lysates (WCLs) were precleared using protein A Sepharose (Sigma-Aldrich) for 30 minutes at 4°C. G6b-B was immunoprecipitated from collagen-WCLs with anti-G6b-B antibody and protein A sepharose overnight at 4°C as previously described (Mazharian et al., 2012).

WCLs were either boiled in SDS-loading buffer and analyzed by SDS-PAGE (NuPage 4-12% Bis-Tris-Gradient Gel) and traditional western blotting or, for quantitative analysis, analyzed with an automated capillary-based immunoassay platform (Wes, ProteinSimple, San Jose, USA), according to the manufacturer’s instructions. Briefly, WCLs were diluted to the required concentration with 0.1X sample buffer, then prepared by adding 5X master mix containing 200 mM dithiothreitol (DTT), 5× sample buffer and fluorescent standards (Standard Pack 1, PS-ST01-8) and boiled for 5 minutes at 95°C. Samples, antibody diluent 2, primary antibodies and anti-rabbit secondary antibody, luminol S-peroxide mix and wash buffer were displaced into Wes 12-230 kDa prefilled microplates, (pre-filled with separation matrix 2, stacking matrix 2, split running buffer 2 and matrix removal buffer, SM-W004). The microplate was centrifuged for 5 minutes at 2500 rpm at room temperature. To start the assays, the capillary cartridge was inserted into the cartridge holder and the microplate placed on the plate holder. To operate Wes and analyze results Compass Software for Simple Western was used (version 3.1.7, ProteinSimple). Separation time was set to 31 minutes, stacking loading time to 21 seconds and sample loading time to 9 seconds. Primary antibodies were incubated for 60 minutes and for detection the High Dynamic Range (HDR) profile was used. For each antibody, a lysate dilution experiment was performed first to confirm the optimal dynamic range of the corresponding protein on Wes. This was followed by an antibody optimization experiment to compare a range of dilutions and select an antibody concentration near to saturation level to allow a quantitative comparison of signals between samples. The optimized antibody dilutions and final lysate concentrations were as follows: anti-Src p-Tyr418 and anti-GAPDH antibodies were used at 1:10 dilution on 0.05 mg/ml lysates; anti-Shp1 p-Tyr564, anti-Shp2 p-Tyr542 and anti-Shp2 p-Tyr580 antibodies were used at 1:10 dilution, and anti-Syk p-Tyr525/6 antibody was used at 1:50 dilution on 0.2 mg/ml lysates.

### Statistical analysis

All data is presented as mean +/− standard deviation (SD). Statistical significance was analyzed by one-way or two-way ANOVA, followed by the appropriate post hoc test, as indicated in the figure legend, using GraphPad Prism 6 (GraphPad Software Inc., San Diego, CA, USA).

## ACKNOWLEDGEMENTS

The authors would like thank all the voluntary blood donors and the phlebotomists as well as Jamie Webster from the University of Birmingham Core Protein Expression Facility, Silke Heising and Louise Tee for excellent technical assistance, Jeremy A. Pike for assistance with image and KNIME analysis, and all members of the Biomedical Services Unit for exceptional maintenance of mouse colonies.

## FUNDING

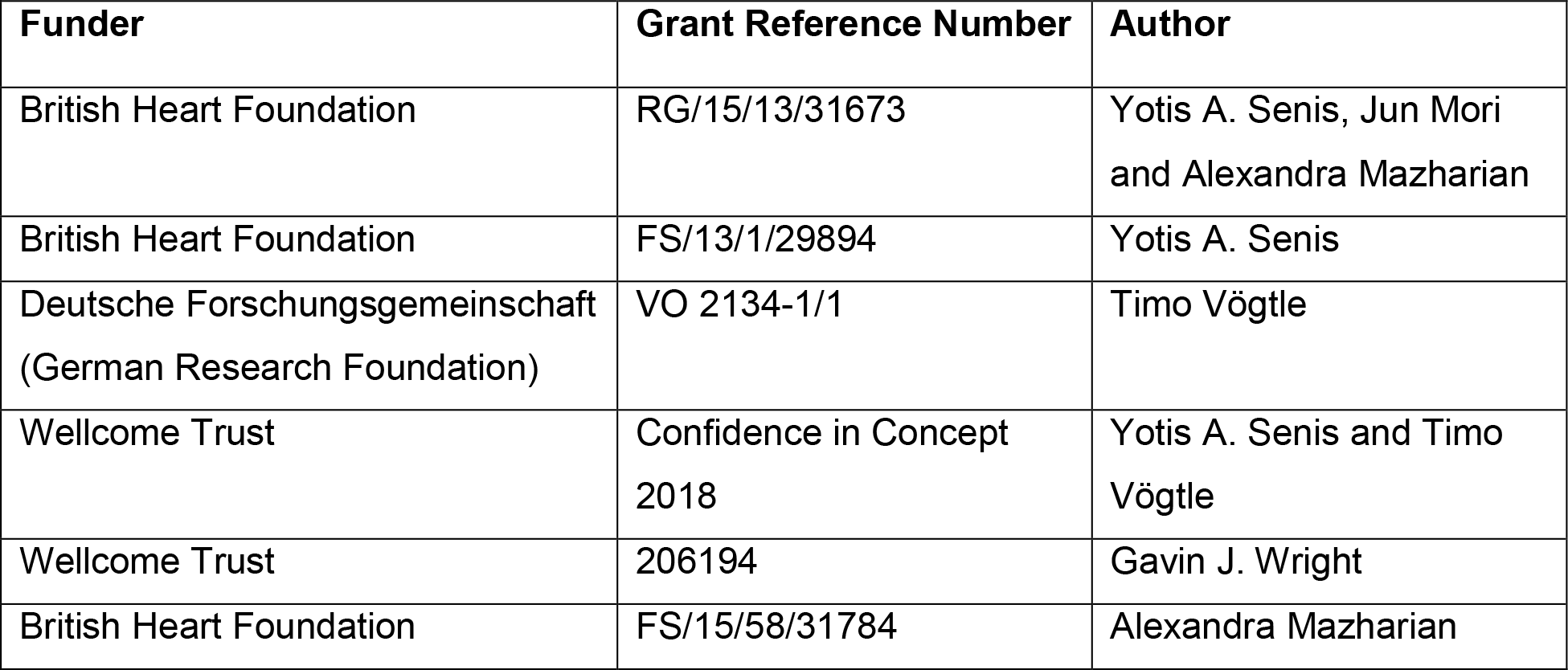

The funders had no role in study design, data collection and interpretation, or the decision to submit the work for publication.

## DATA AVAILABILITY

Atomic coordinates have been deposited with the PDB with the accession numbers: 6R0X. The following datasets were generated:

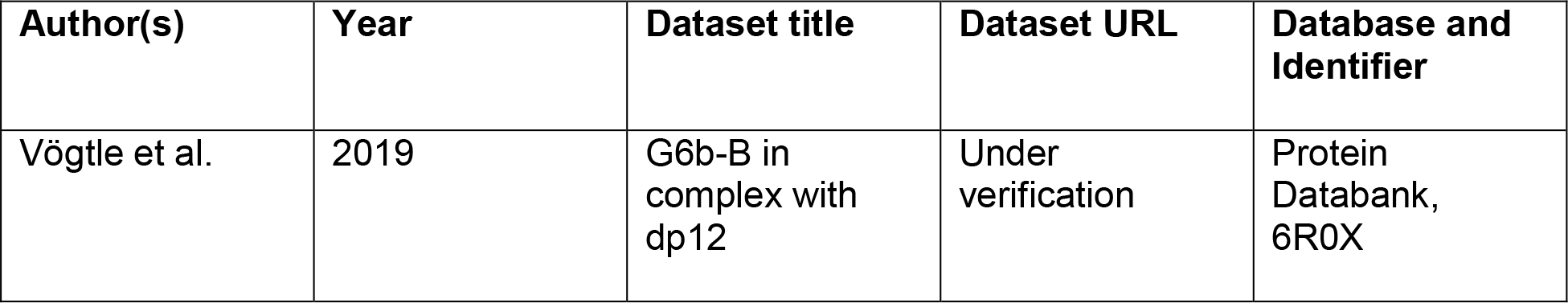

| Author(s) | Year | Dataset title | Dataset URL | Database and Identifier |
| --- | --- | --- | --- | --- |
| Vögtle et al. | 2019 | G6b-B in complex with dp12 | Under verification | Protein Databank, 6R0X |

---

**Figure 2–figure supplement 1.**
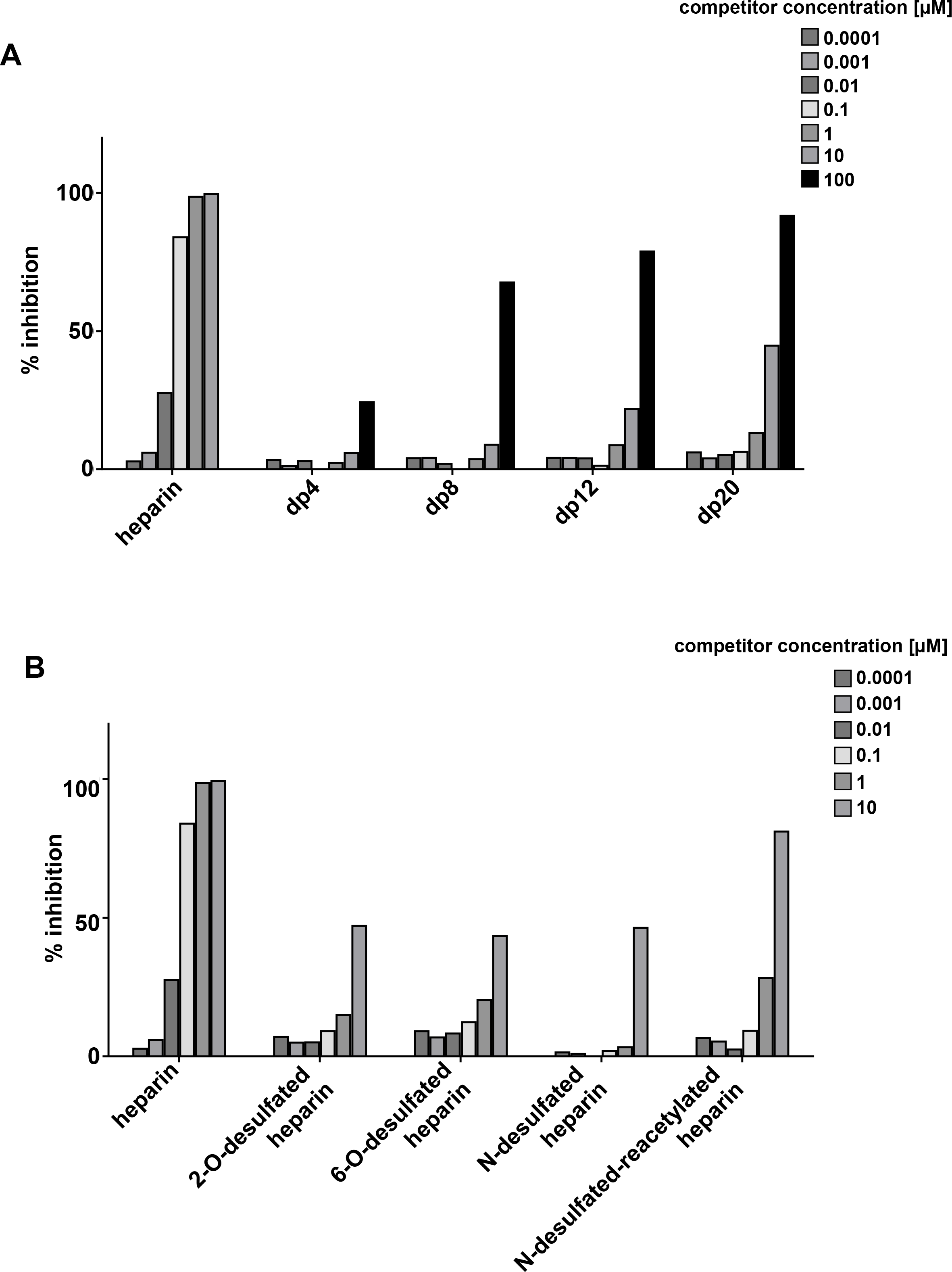
Loss of heparin sulfation impairs interaction with G6b-B. 96-well plates were coated with streptavidin followed by incubation of biotinylated heparin and binding huG6b-B monomer/ anti-G6b-B antibody complex (1 μg/ml each) was measured in the presence of **(A)** heparin or oligosaccharide defined length (dp, degree of polymerization = number of saccharides) or **(B)** selectively desulfated heparin species at the given concentration. % inhibition was calculated by normalizing OD to maximum and minimum values. Representative results for 2 independent experiments.

**Figure 4–figure supplement 1.**
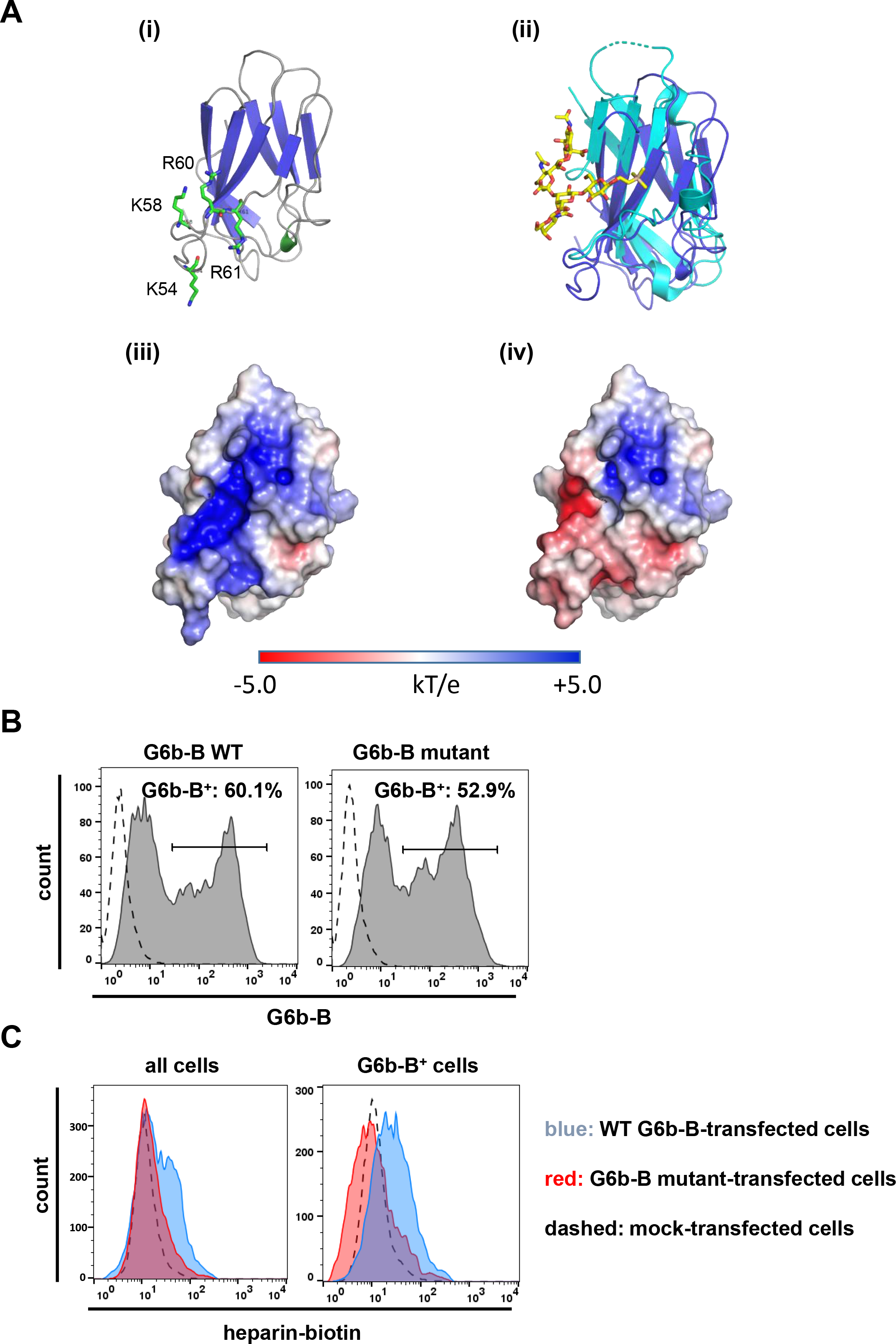
Mutations in G6b-B abolishes heparin binding. **(A)** G6b-B model (i) ribbons representation showing predicted locations of basic residues K54, K58, R60 & R61 in green (ii) superposition of G6b-B model (dark blue) with Siglec-7 (cyan) in complex with its sialic acid ligand (yellow) (iii) & (iv) electrostatic surface potentials as calculated by ABPS for both the wild-type G6b-B model and the G6b-B K54D, K58D, R60E, R61E mutant respectively. Positively charged surfaces are blue and negatively charged are red. **(B, C)** CHO cells were transiently transfected with wild-type (WT) or mutant G6b-B and analyzed by flow cytometry. **(B)** Staining of cells with an anti-G6b-B antibody confirms similar transfection efficacy and G6b-B expression. **(C)** CHO cells were incubated with biotinylated heparin, followed by PE-labelled streptavidin. Heparin-biotin binding of all cells (left) and G6b-B^+^ cells (right), was measured. Representative results from two independent experiments.

**Figure 4–figure supplement 2.**
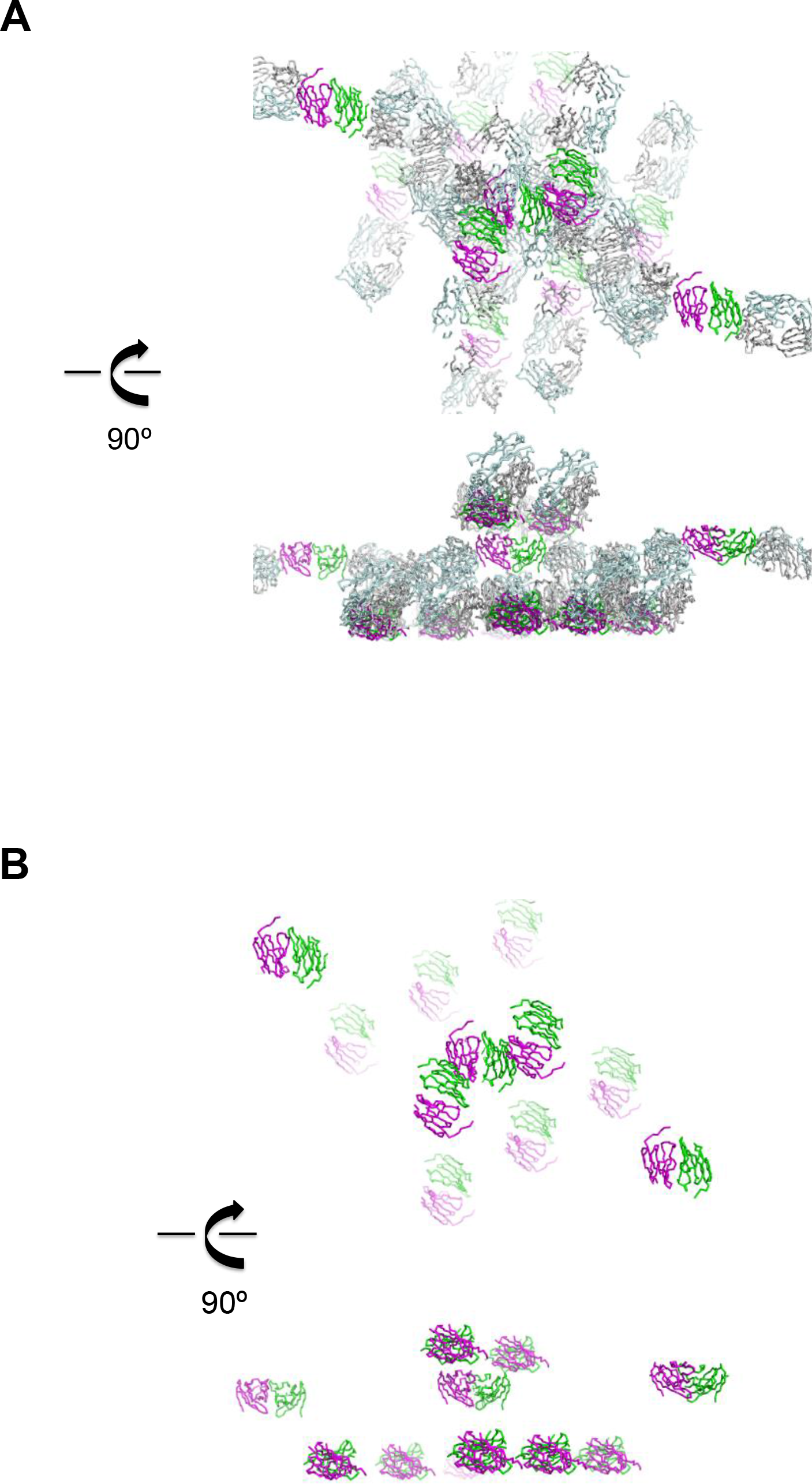
Representation of the crystal lattice. **(A)** Crystal lattice of the ternary complex of G6b-B bound to heparin and the G6b-B specific Fab fragment. Top and bottom half the view are related by a 90º rotation about the horizontal axis. Proteins are shown as Cα traces, with G6b-B in magenta and dark green, and the Fab fragment chains in light blue and light grey, respectively. **(B)** Illustration of crystal lattice contacts by the G6b-B ECD dimer. The view shown is the same as in (A), but omitting the Fab fragment chains. Top and bottom half the view are related by a 90º rotation about the horizontal axis.

**Figure 4–figure supplement 3.**
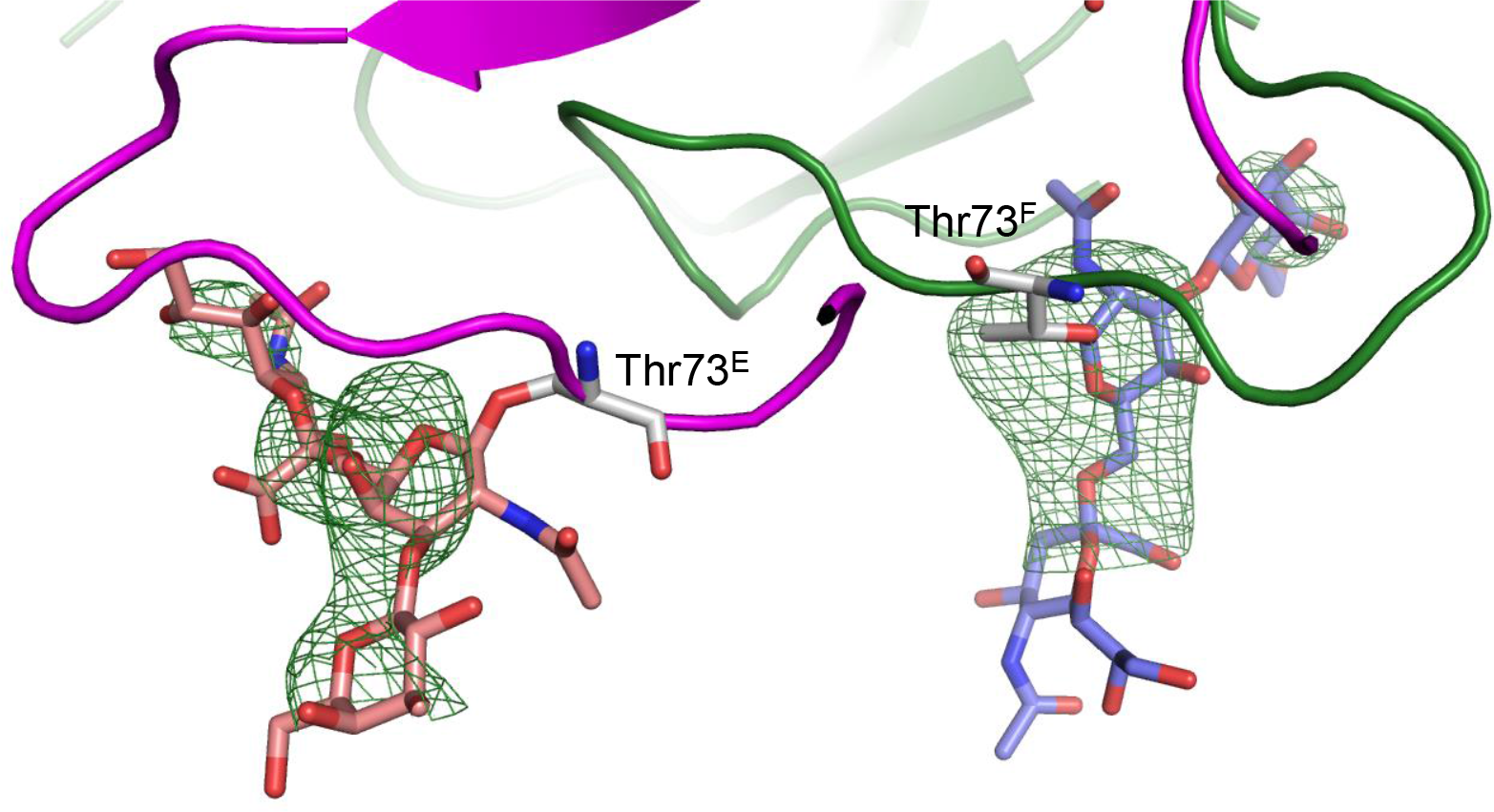
Unbiased σA-weighted difference density map demonstrating the presence of the O-linked glycosyl groups at Thr73.

**Figure 6–figure supplement 1.**
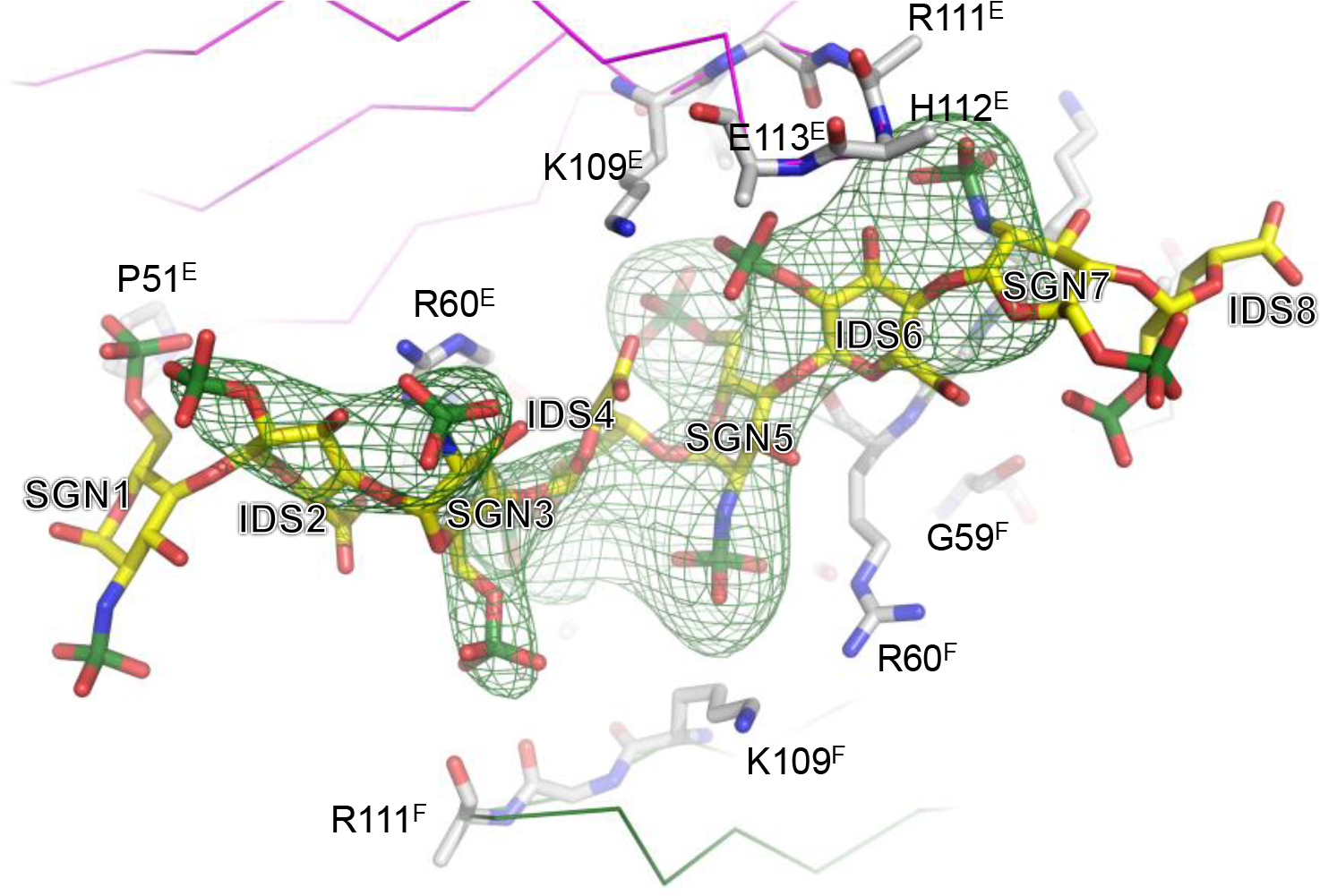
Unbiased σ_A_-weighted difference density map demonstrating the presence of the heparin ligand. The electron map has been calculated with phases of from a structure model lacking the heparin atoms and with amplitudes (Fo – Fc), contouring the map at 2.8 σ above the mean.

**Figure 11–figure supplement 1.**
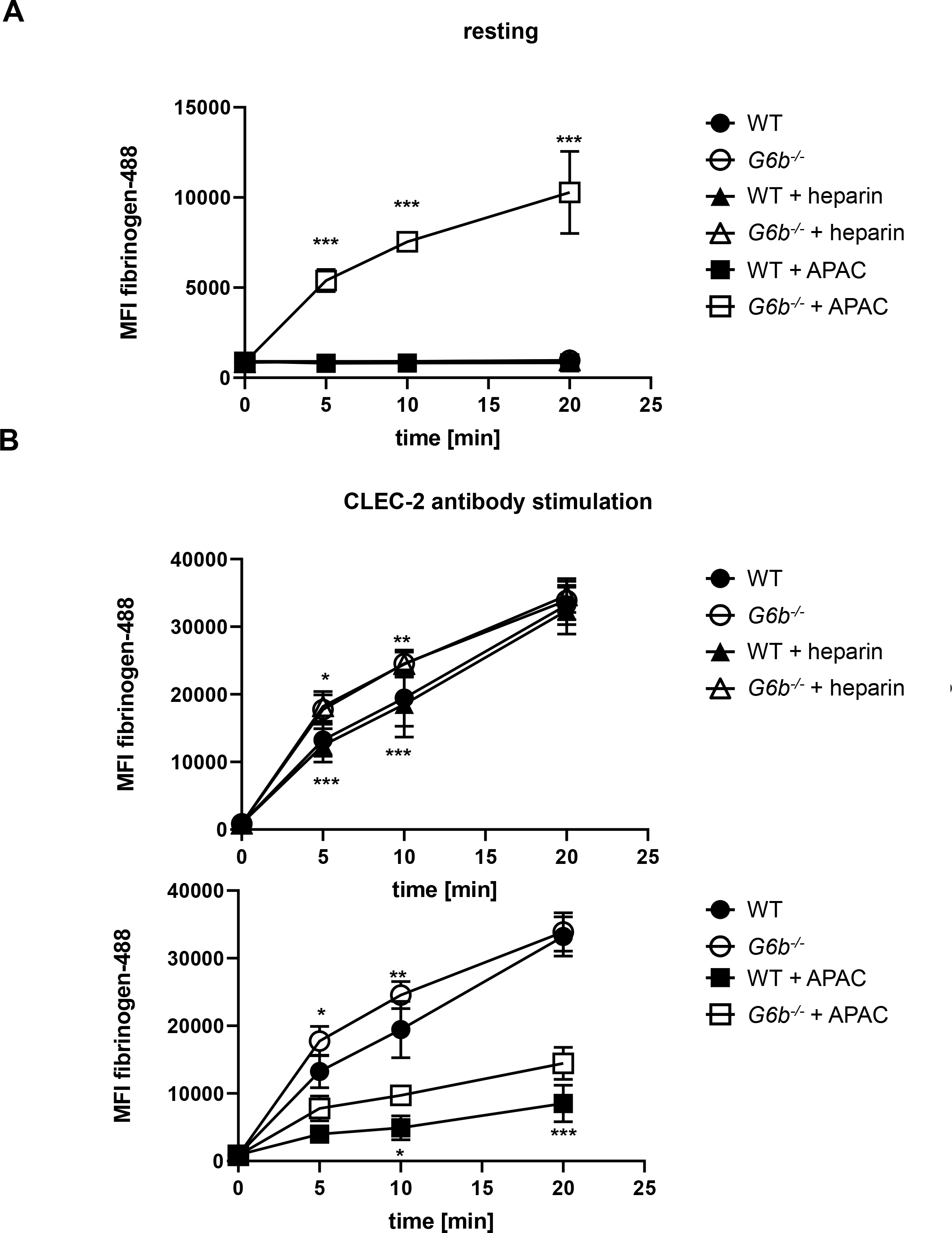
Effect of heparin and APAC on fibrinogen binding of WT and *G6b* KO platelets. Mouse blood was incubated with the indicated compounds (0.05 μM) in the **(A)** absence or **(B)** presence or of a stimulating CLEC-2 (3 μg/ml) for the indicated time. Samples were fixed and fibrinogen-Alexa488 binding, a measure of integrin activation, was determined by flow cytometry. n=5-6 mice/condition/genotype from 2 independent experiments. P-values were calculated using (A) two-way ANOVA with Sidak’s post-hoc test (comparison of WT APAC vs *G6b*^−/−^ APAC) or (B) two-way ANOVA with Turkey’s post-hoc test and refer to the difference between WT and *G6b*^−/−^ ***, *P*<0.001, **, *P*<0.01 and *, *P*<0.05

**Figure 11–figure supplement 2.**
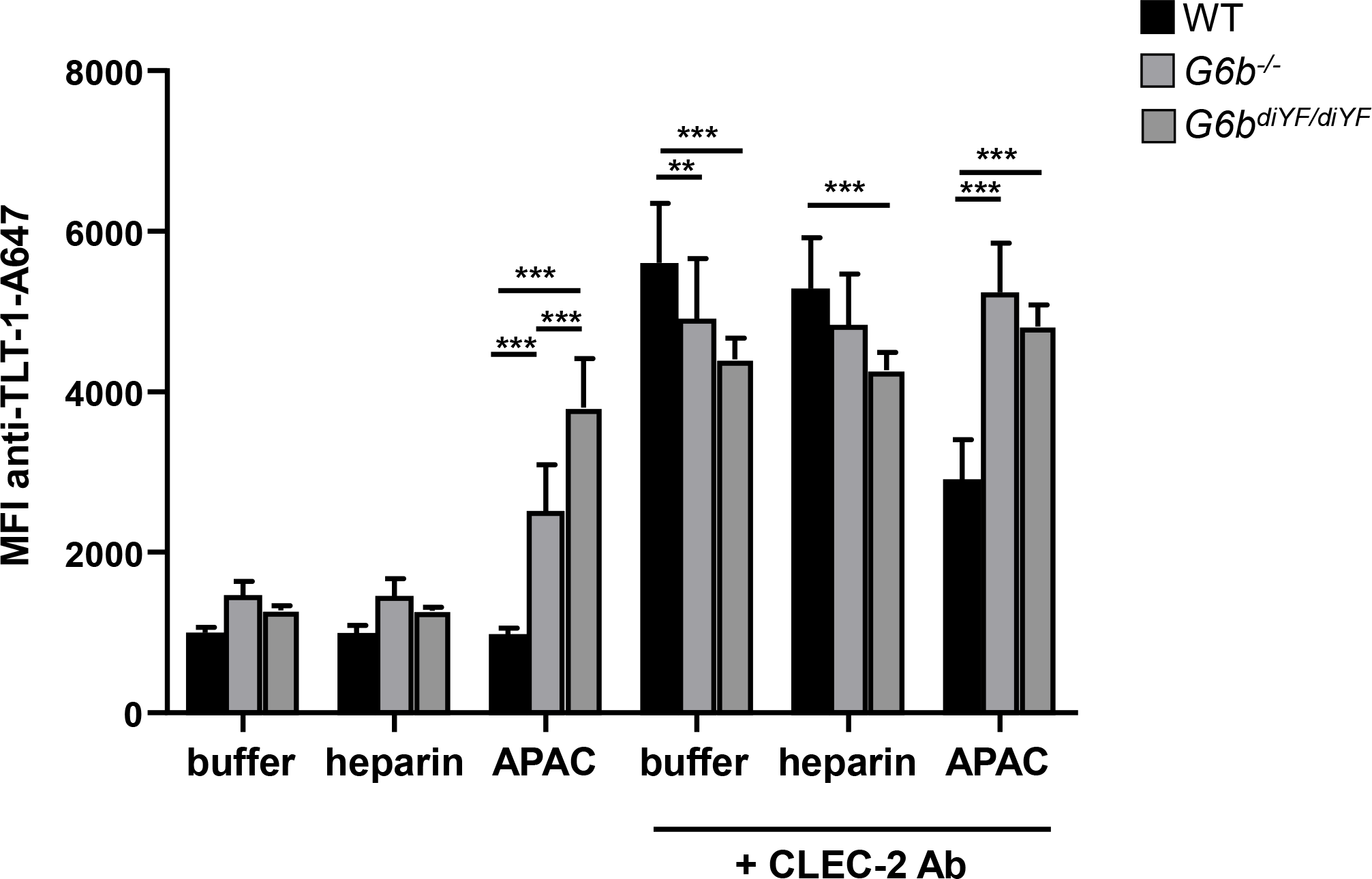
G6b-B signaling is required for the inhibitory effect of APAC on platelet degranulation. Mouse blood was incubated with the indicated compounds for 20 min. Samples were fixed and TLT-1 surface levels were determined by flow cytometry. n=6-8 mice/condition/genotype from 3 independent experiments. P-values were calculated using two-way ANOVA with Turkey’s post-hoc test ***, *P*<0.001; and ** *P*<0.01.

**Figure 12–figure supplement 1.**
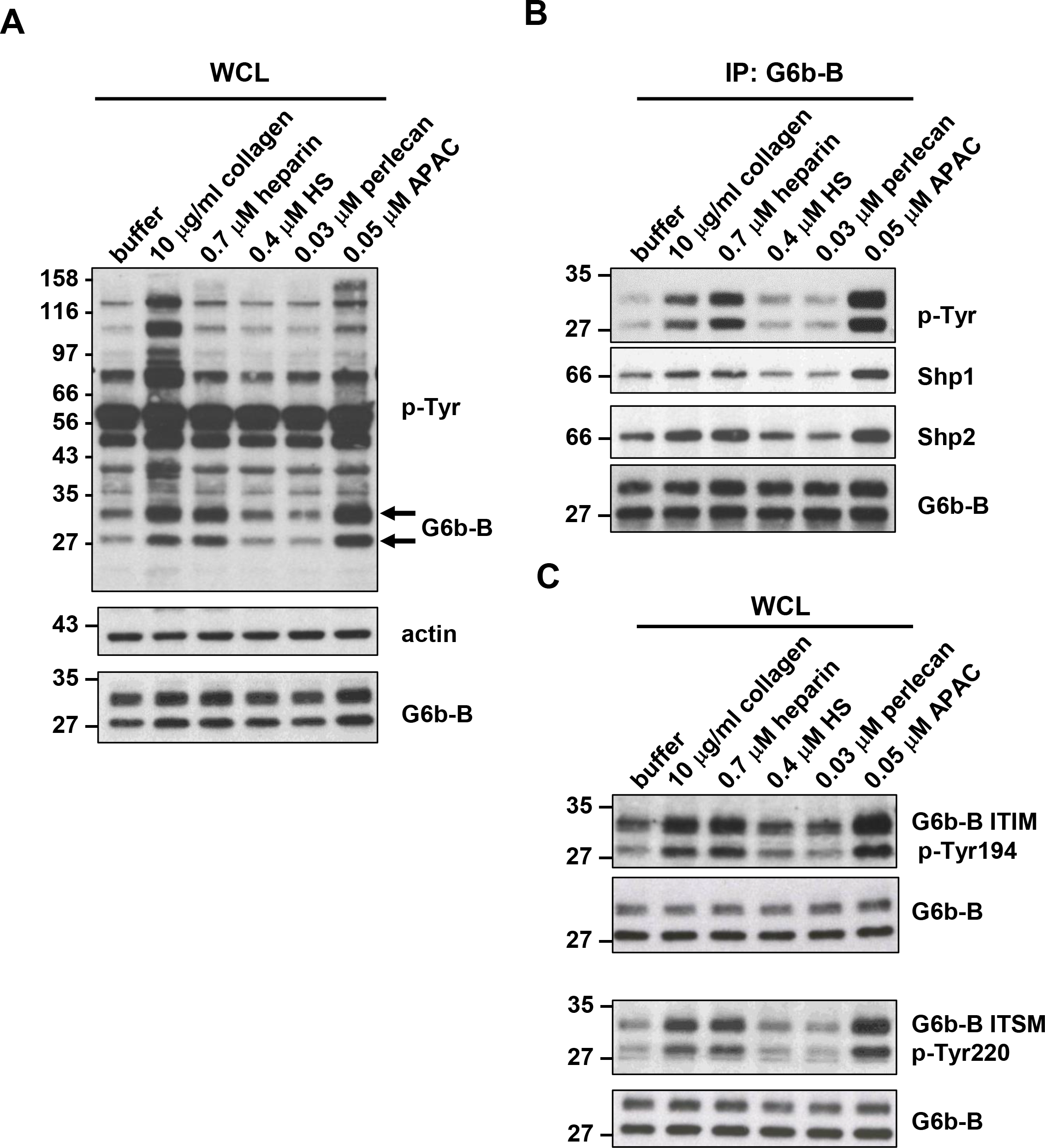
Effects of G6b-B ligands on G6b-B phosphorylation. Washed human platelets (5×10^8^/ml) were incubated for 90 s with the indicated compounds in the presence of 10 μM integrillin. Whole cell lysates (WCL) were directly analyzed by western blotting **(A, C)** with the indicated antibodies or **(B)** subjected to immunoprecipitation (IP) with an anti-human G6b-B antibody, followed by western blotting. Representative results of 3 independent experiments.

**Figure 12–figure supplement 2.**
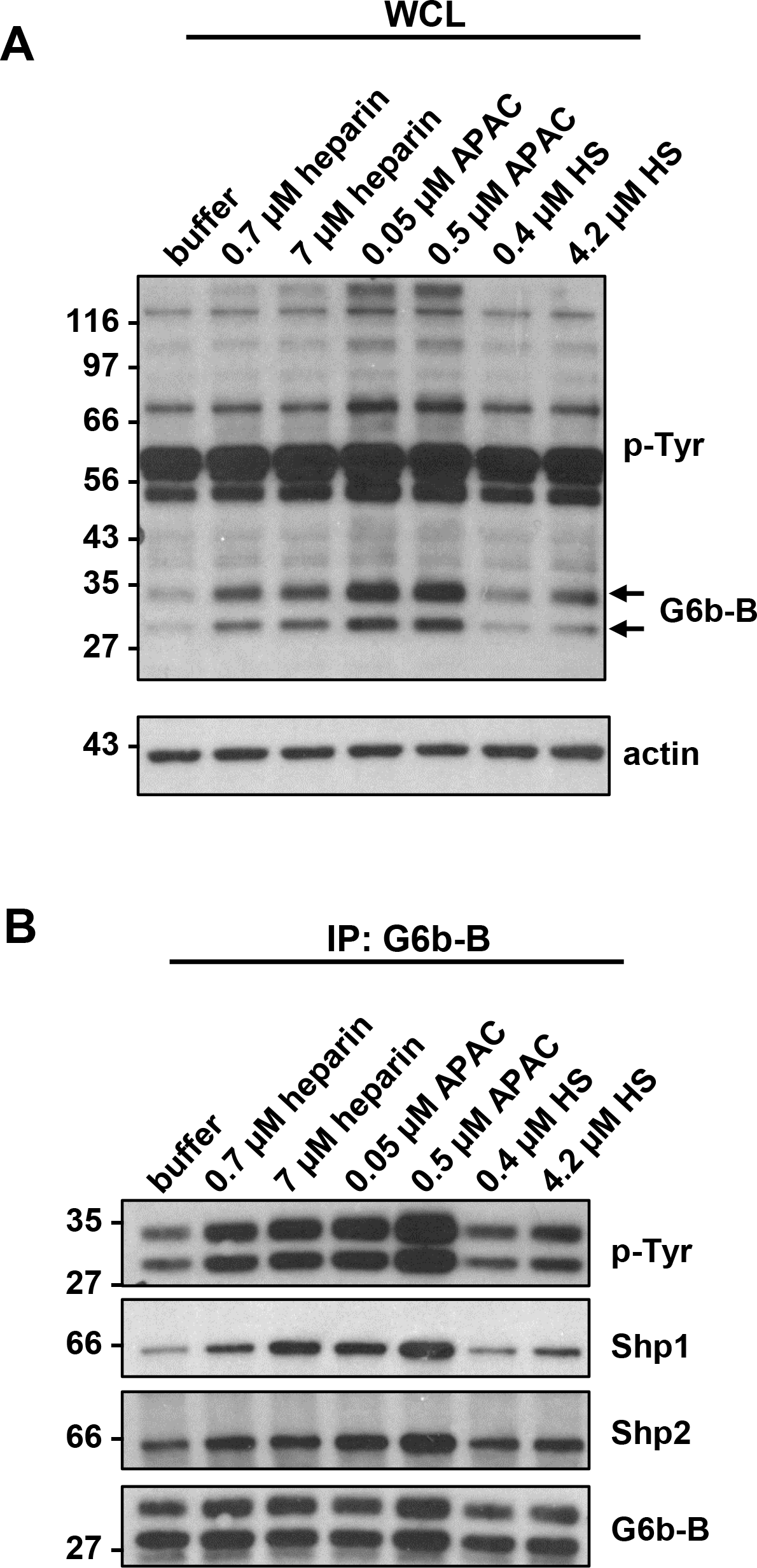
Effects of high doses of G6b-B ligands on G6b-B phosphorylation. **(A, B)** Washed human platelets (5 × 10^8^/ml) were incubated for 90 s with the given compounds at the given concentration in the presence of 10 μM integrilin. Samples were lysed and whole cell lysates (WCL) were directly analyzed by western blotting **(A)** with the indicated antibodies or **(B)** subjected to immunoprecipitation (IP) with an anti-human G6b-B antibody, followed by western blotting.

## Supplementary files

**Supplementary Figure 1.**
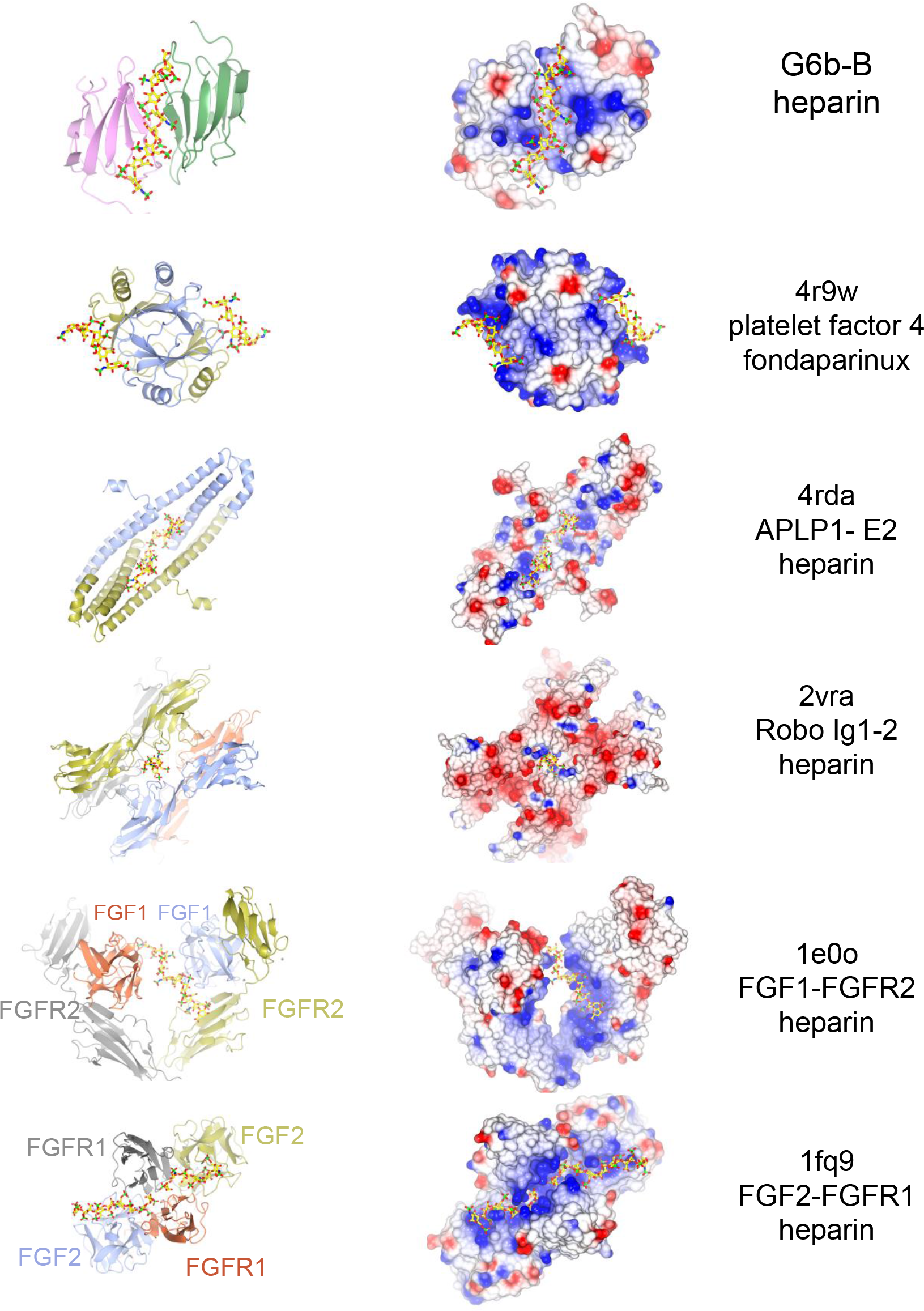
Selected structures of proteins with a heparin ligand. Side-by-side views of Cα traces (ribbon representation) and electrostatic surfaces of proteins bound to heparin or a heparin analogue. The subset includes structures where the heparin ligand bridges two or more subunits. At the time of writing, the PDB contained 33 proteins structures with heparin as a ligand, in addition to a few structures with heparin analogues. From top to bottom: G6b-B (Fab fragments and glycosyl chains omitted from view); 4r9w – platelet factor 4 bound to fondaparinux (synthetic heparin analogue) (Cai et al., 2015); 4rda – E2 domain of amyloid precursor protein-like protein 1 (APLP1) (Dahms, Mayer, Roeser, Multhaup, & Than, 2015); 2vra - immunoglobulin-like domains 1 and 2 of Drosophila Robo (Fukuhara, Howitt, Hussain, & Hohenester, 2008); 1e0o – ternary complex of FGF1, FGFR2 and heparin (Pellegrini, Burke, von Delft, Mulloy, & Blundell, 2000); 1fq9 - ternary complex of FGF2, FGFR1 and heparin (Schlessinger et al., 2000). The coloring of the electrostatic surfaces a potential scale from −0.5 V (−19.5 kT/e, red) to +0.5 V (+19.5 kT/e, blue). The ribbon representations are colored by peptide chain, and heparin is shown as a stick model with color codes: C – yellow, O – red, N – blue, S – green.

